# Diverse ways of holding verbal information in mind revealed with functional Magnetic Resonance Imaging and Transcranial Magnetic Stimulation: Individual differences in left anterior parietal cortex

**DOI:** 10.1101/2025.10.30.685124

**Authors:** Nicola J. Savill, Piers Cornelissen, Meichao Zhang, André Gouws, Jonathan Smallwood, Elizabeth Jefferies

## Abstract

Cognitive neuroscience has identified brain systems that reliably underpin specific abilities, including verbal short-term memory. However, there is no simple mapping between anatomy and function; moreover, brains are differently organised across people. In two functional magnetic resonance imaging studies, and using transcranial magnetic brain stimulation to test causality, we characterised differences in the neural processes supporting verbal short-term memory in healthy participants who varied in their use of semantic information. In group analyses, left anterior inferior parietal cortex showed the expected pattern for a “phonological buffer” – this site was responsive to phonological tasks and verbal short-term memory for meaningless items. However, the functions of this site differed across the sample; individuals with the strongest “semantic reliance” in short-term memory (showing higher imageability effects and poorer nonword performance) showed weaker phonological responses. They also showed more activation when maintaining meaningful word sequences in short-term memory, compared with nonwords. In these semantically-reliant participants, left anterior inferior parietal cortex showed stronger functional connectivity to limbic and default mode regions during nonword rehearsal. While inhibitory stimulation to phonologically responsive regions disrupted nonwords more than words in people with good phonological short-term memory, these effects were reversed when verbal short-term memory was more semantically reliant. These findings show that left anterior inferior parietal cortex supports the maintenance of different kinds of verbal information across individuals.

## Introduction

Keeping verbal information in mind is a critical component of cognition, underpinning thought, communication, learning, and arithmetic (Baddeley, 2003). Verbal short-term memory (STM) is supported by the buffering of phonological information in dorsal language regions, including left parietal cortex encompassing the supramarginal gyrus (Jacquemot and Scott, 2006; Majerus, 2013). This site is consistently implicated in phonological processing and maintenance (Buchsbaum and D’Esposito, 2008; Papagno et al., 2017). However, a key question is whether its functional organisation can differ across individuals, reflecting variability in how verbal information is maintained and processed. This question is important for neurocognitive models of verbal short-term memory, which seek to explain how phonological and semantic processes interact to support flexible verbal maintenance.

Word knowledge contributes to verbal STM, as lexical-semantic information can reconstruct incomplete phonological traces via interactive activation between phonological and semantic representations during rehearsal (Jefferies et al., 2006b; Acheson and MacDonald, 2009). This explains why sequences of meaningful words are maintained better than nonwords (Brener, 1940; Hulme et al., 1991; Savill et al., 2019a). Since phonological and semantic representations mutually constrain one another, phonological weakness can be compensated by semantic recruitment and vice versa (Patterson and Lambon Ralph, 1999; Foygel and Dell, 2000). While STM also draws on cognitive control, retention of meaningful sequences is thought to largely depend on automatic activation of long-term semantic knowledge (Jefferies et al., 2004; Schwering and MacDonald, 2020), which can stabilise fragile phonological representations in STM (Patterson et al., 1994; Jefferies et al., 2006a, 2006b).

Phonological and semantic processes rely on dissociable neural mechanisms, which can vary independently across individuals. The left supramarginal gyrus (SMG) supports phonological buffering and is particularly responsive to nonwords and sequence length, whereas the left anterior temporal lobe (ATL) supports long-term semantic representations and is sensitive to lexical-semantic factors such as imageability (Hoffman et al., 2015; Savill et al., 2019a). Neuropsychological studies demonstrate a reciprocal pattern: individuals with phonological impairments rely more heavily on semantic knowledge, while those with semantic impairments depend more on phonological processing (Jefferies et al., 2008; Verhaegen et al., 2013). Neuroimaging and TMS studies in healthy participants converge on the same principle, showing that those who rely more on word meaning recruit the ATL more strongly during reading and recall (Hoffman et al., 2015; Woollams et al., 2017). These findings suggest that the balance between semantic and phonological mechanisms is not fixed; but varies across individuals in ways that shape language performance.

In a behavioural study of over 80 participants, we observed a common division of labour across phonology and semantics: participants with weaker phonological skills showed stronger effects of imageability in reading and immediate serial recall (Savill et al., 2019b). Building on this, the current study used fMRI and TMS to characterise differences in neural organisation supporting verbal STM in participants with strong and weak “semantic reliance”: participants, recruited from our prior sample, were considered to be more semantically reliant when they showed higher imageability effects and poorer nonword performance, suggesting greater involvement of conceptual processing or imagery, and weaker capacity to maintain phonological information in the absence of meaning. Here, they completed phonological and semantic tasks (Experiment 1) and verbal STM tasks involving nonwords, random words, and meaningful sequences (Experiment 2). Inhibitory TMS was delivered to individually defined SMG and ATL sites to test the causal role of these regions in participants differing in semantic reliance (Experiment 3).

These experiments allow us to evaluate accounts of the brain basis of individual differences in verbal STM (Hartwigsen et al., 2017). One view predicts that semantic-reliant individuals recruit semantic regions more during immediate recall. An alternative view is that individual differences reflect remapping of function across cortex, such that the same region can support different aspects of cognition to varying degrees. Within this framework, classical STM regions in inferior parietal cortex like SMG may buffer diverse kinds of information, supported by variable connectivity patterns across the brain.

## Materials and Methods

### Participants

The study was approved by the ethics boards of the Department of Psychology and York Neuroimaging Centre at the University of York. Participants were 30 healthy, right-handed native English adults with normal vision and hearing and no reported history of learning difficulties or developmental, neurological or psychiatric disorders screened for TMS and MRI safety. These participants were specifically invited to take part on the basis of their semantic reliance in auditory-verbal short-term memory as tested in an earlier study (Savill et al., 2019b): the screening task used to identify candidate participants for this follow-up study tested immediate serial recall (ISR) of high- and low-imageability word lists (six and seven items) and nonword lists (four and five items). Fifteen participants showing large effects of word imageability (as an indirect measure of semantic support) were identified for comparison with another group of fifteen who showed little to no impact of imageability on recall (such that their recall differentially responded to imageability, at the group level, by a very large effect size; Cohen’s d = 2.68). Importantly, these participants at relative behavioural extremes in terms of stimuli effects in ISR were identified after also ensuring at-least average standardised reading achievement, similar word reading fluency, nonverbal IQ, forward digit span, and semantic knowledge (see Supplementary Table S1). In line with the association between phonological function and imageability effects observed across different language tasks by Savill et al. (2019b), these groups differed in terms of their phonological capacity: individuals selected on the basis of their larger imageability effects tended to have poorer recall of nonwords and low-imageability words than those with small-to-no effects of imageability on recall performance. Since participants showing strong semantic effects and weak phonological retention are more reliant on semantic information in short-term memory, these groups are referred to as high and low semantic reliance (SR) below (shown in red and blue in Figure 1).

**Figure 1.**
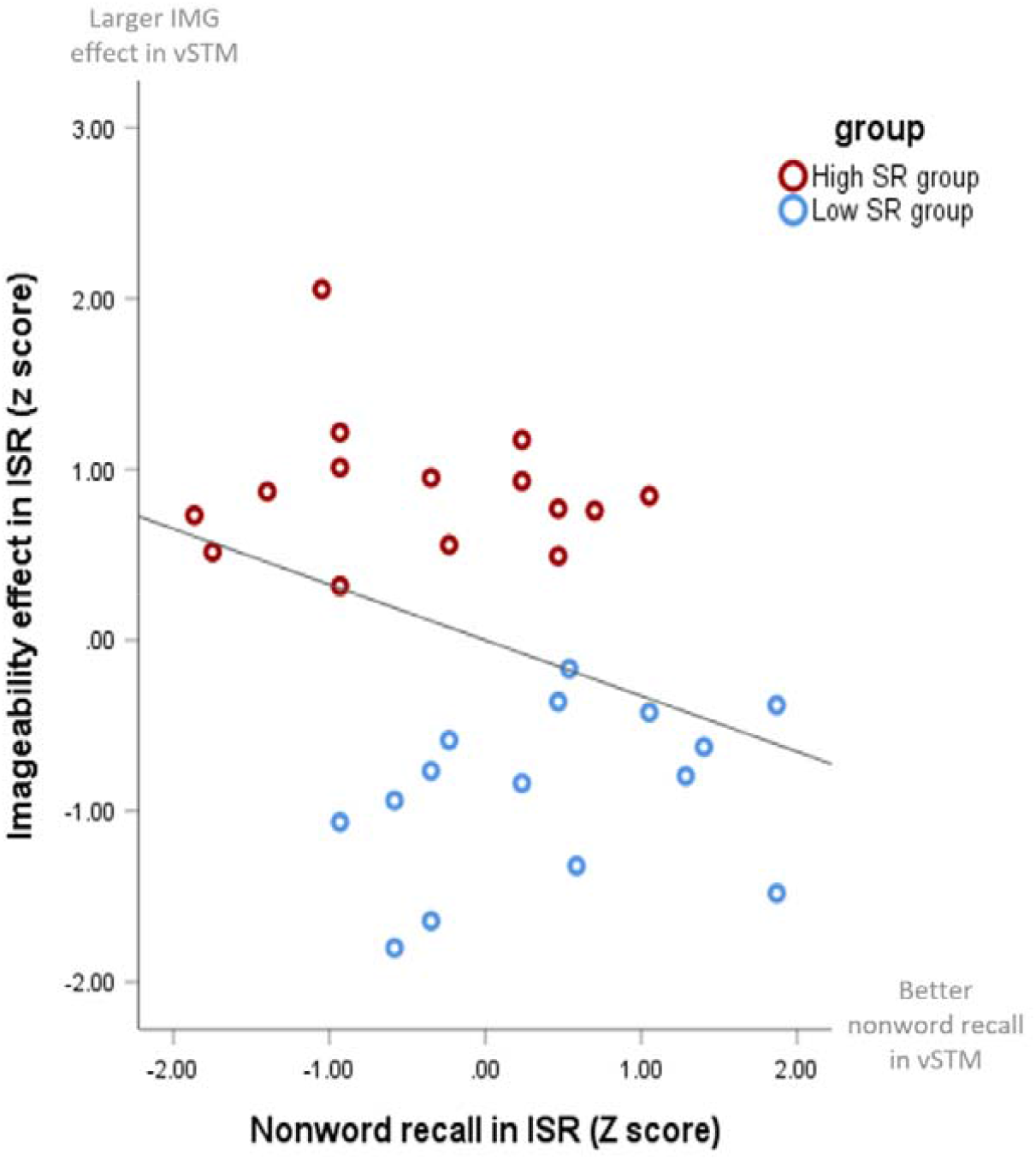
The behavioural relationship between the size of imageability effect and nonword performance in immediate serial recall in participants selected for the present study on the basis of relatively large imageability effects (high semantic reliance) or minimal imageability effects (low semantic reliance) in this task outside the scanner. The imageability effect was computed by dividing the score for low imageability word recall by the score for high imageability word recall and calculating inverse z scores of those values. A larger value on the y axis denotes a larger imageability effect (i.e., poorer recall of low imageability words relative to high imageability words). Participants who showed little performance detriment from the presentation of low imageability words (low semantic reliance group) also tended to have better nonword recall. SR = semantic reliance. ISR = immediate serial recall. IMG = imageability. vSTM = verbal short-term memory.

All 30 participants completed the fMRI component of the study, and 25 of these (13 high and 12 low semantic reliance) completed the TMS component. Participants were compensated for their participation at a rate of £10 an hour. All participants gave written consent prior to their participation.

#### Measurement of semantic reliance in participants

To account for the correlation between imageability effects and nonword performance in immediate serial recall, shown for our sample in Figure 1, we followed the approach in our previous behavioural study and computed a composite measure combining these aspects of performance (Savill et al., 2019b). For each participant we calculated the difference between their inverse z-scored imageability effect and z-scored nonword performance in the immediate serial recall screening task to generate a single behavioural measure of semantic reliance. This score was used as a regressor for fMRI analyses to avoid problems of collinearity, since participants’ use of semantic versus phonological information in immediate serial recall are correlated and inter-dependent (comparison with an alternative approach of combining these scores in provided in Supplementary Materials, Section C: the two approaches generated near-identical scores). The highest semantic reliance values corresponded to relatively large imageability effects (i.e., better recall of high than low imageability words) and poor nonword recall in ISR. The lowest semantic reliance values corresponded to small imageability effects and relatively strong nonword recall performance (Table S1 in Supplementary Materials).

### In-scanner experimental tasks

#### fMRI Task 1: Phonological, semantic and case judgements

We scanned the participants while they categorised 96 words according to their phonological and semantic properties (see Figure 2). With a two-alternative forced choice button press (Yes/No), participants indicated whether each word, singly presented in the centre of the screen, consisted of two syllables (phonological judgement task), or whether it represented a natural object (semantic judgement task). In a third low-level condition, designed to provide a visuo-motor baseline for these language tasks, participants determined if unpronounceable consonant strings were presented in upper or lower case.

**Figure 2.**
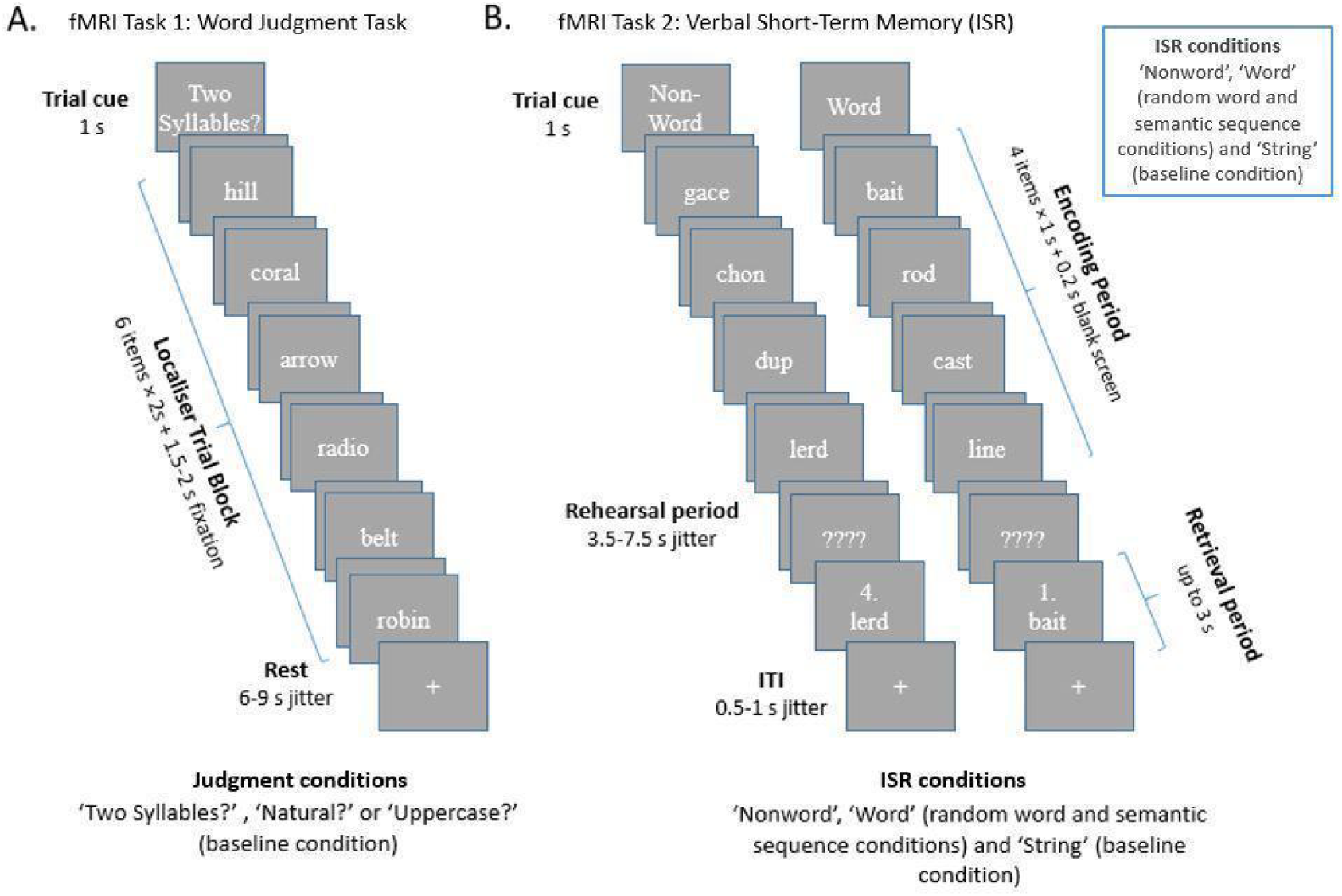
In-scanner experimental tasks. (A) Example of a word judgement task. The cue (‘Two Syllables?’, ‘Natural?’, ‘Uppercase?’) determined whether the block required phonological, semantic or case decisions. Event-related analyses assessed functional responses to each trial item. (B) Example nonword and semantic sequence trials in the verbal short-term memory task. A third condition presented unrelated random words and a fourth visual baseline condition presented strings of the same capital letter (e.g., AAAA × 4). Participants were asked to maintain the series of four items and were tested on their recall a few seconds later (by asking them to decide if a probe was presented in a particular list position). The trial cue specified whether letter strings, nonwords or words would be presented.

#### Stimuli for phonological, semantic and case judgements

Ninety-six mono-, bi-and tri-syllabic words were selected from available imageability norms (Bennett et al., 2011; Cortese & Fugett, 2004; Schock, et al., 2012). The words were selected so that they could be used in both semantic and phonological decision conditions (average lexical frequency, M = 4.07, SD = 0.48 (van Heuven et al., 2014); imageability, M = 6.34, SD = 0.30; word length, M = 5.77, SD = 1.21). There were equal numbers of YES and NO responses within each decision. Consonant strings for the baseline task matched the number of letters in the words and the letters were selected so that the ascender height of the lowercase form was the same height as its uppercase form, e.g., HHFFHL vs. hhffhl). Each stimulus was presented in a white Times New Roman font on a grey background. Task stimuli can be found at https://osf.io/s68q9/.

Categorisation performance was piloted in a sample that did not complete the main experiment (N=10; mean age = 30.9 years; 6 females). This revealed no significant differences between the three tasks in reaction time or accuracy (see Supplemental Materials Section A).

#### Experimental procedure for phonological, semantic and case judgements

An example of the trial structure can be seen in Figure 2. The three tasks (phonological, semantic, case judgements) were presented in blocks of 6 trials. Each block started with an instruction screen for 1s (‘Natural?’, ‘Two Syllables?’ or ‘Upper Case?’). Each item was then presented for up to 2s; the trial item was replaced with a fixation cross following a response for the remainder of this period. The fixation cross remained for an additional jittered inter-trial interval, lasting 0.5 – 1s. Participants responded via a button response box in their left hand, using the index and middle fingers to respond yes or no.

Participants had a maximum of 3s from item onset to respond. These six-item blocks (approximately 16s) were separated either by a jittered fixation cross lasting 6 – 9s, after which 6 more trials from the same task were presented (e.g., phonological block – null event – phonological block), or a jittered rest period of between 8 – 10s, with a green fixation cross indicating a switch between tasks. There were two experimental runs, with 8 blocks per judgment condition in each run (i.e., 16 blocks per judgment condition across runs; 24 blocks in total per run; judgment conditions switched every 4 blocks). Within each run, each word was presented once in the phonological or semantic judgment task, and blocks were presented in one of two fixed pseudorandom orders, starting with either the semantic or phonological judgement task (counterbalanced across participants). Stimulus presentation and response recording were controlled using PsychoPy (Peirce, 2009). Psychopy script available here: https://osf.io/s68q9/.

#### fMRI Task 2: Immediate serial recall

An immediate serial recall (ISR) task was used to measure brain activity during the verbal maintenance of meaningful and meaningless materials (Figure 2). This task differed from the behavioural screening task in several ways: list length was set at four items for all conditions to minimise errors (since our focus was on short-term memory activation during successful trials), items were presented visually rather than auditorily (to minimise the impact of concurrent scanner noise on performance) and verbal recall was assessed with a button press instead of a spoken response (to minimise movement associated with articulating a spoken response). Four conditions were tested: (1) a simple visual baseline, consisting of strings of the same uppercase letter (e.g., BBBB, BBBB, BBBB, BBBB); (2) nonword lists of four pronounceable nonwords, used to assess phonological maintenance independent of meaning and familiarity (e.g., gace, chon, dup, lerd; information on stimuli provided below); (3) word lists included four random unrelated words (e.g., zone, book, jam, cloud) and (4) semantic sequences presented a series of words that formed a coherent, meaningful sequence (e.g. bait, rod, cast, line). In all four conditions, a few seconds after the end of the list, participants were probed with an item and a location in the list (1^st^, 2^nd^, 3^rd^ or 4^th^) and were asked to decide whether the probed item was presented in that position (pressing one of two buttons to indicate a YES/NO judgement, using the index and middle fingers of their left hand).

#### Stimuli for immediate serial recall

Lists comprised written monosyllabic words or pronounceable nonwords, chosen to avoid phonological overlap between items within a list (i.e., no phoneme was repeated in a given syllabic position within a list) and repetition of stimuli across the whole set. Meaningful sequences of four nouns (including nouns that could act as multiple parts of speech, e.g., bait, rod, cast, line) were constructed and the 48 that received the highest average semantic coherence ratings in an independent pilot sample of 10 participants were selected. Random word lists were constructed so that average lexical frequency (Sequences, M = 4.54, SD = 0.34; Random Words, M = 4.51, SD = 0.28), word length (Sequences, M = 4.18, SD = 0.48; Random Words, M = 4.17, SD = 0.34), imageability ratings (Sequences, M = 5.42, SD = 0.63; Random Words, M = 5.45, SD = 0.59) and numbers of singular versus plural forms were matched across word conditions (i.e. random word and sequences conditions). Stimuli were presented in a white Times New Roman font on a grey background. Stimuli and Psychopy script here: https://osf.io/s68q9/.

#### Experimental procedure for immediate serial recall

In a given trial, participants first saw a prompt lasting 1s indicating whether letter strings, nonwords or words would be presented. This was followed by the four items to be maintained, presented at a rate of 1/s. Each item was on screen for 0.8s followed by a blank screen for 0.2s. Next, to provide an extended stimulus-independent rehearsal period for analysis, a string of four question marks (‘????’) was presented for a randomly-jittered duration of between 3.5 – 7.5s; participants were instructed to mentally rehearse the sequence in preparation for a test probe. Recall was tested by presenting a probe item in the centre of the screen together with a number (denoting the 1st, 2nd, 3rd, or 4th position in the list). Participants indicated via a button-press whether the probe item was present at that position in the sequence. Half of the ISR trials required a NO response; these could contain a trial item in the wrong position or a new item combining phonemes from items in the list (e.g., in the example semantic sequence trial, ‘1. bait’ would require a YES response, whereas ‘1. rod’ or ‘1. rate’ would require NO responses). There were no incorrect responses for the visual baseline. Participants responded via a button response box in their left hand, using the index and middle fingers to respond yes or no. Participants had a maximum of 3s to respond.

Trials were separated by a jittered inter-trial interval in which a white fixation cross was presented for 0.5 – 1s. After every four trials, the jittered inter-trial interval was replaced by 12s rest (indicated by a green fixation cross). Each experimental condition had 24 trials, divided equally across three runs (8 trials per condition per run; 96 trials in total). These trials were presented in a fixed mixed pseudo-random order within each run and the order of runs was counterbalanced across participants. Stimulus presentation and response recording were controlled using PsychoPy (Peirce, 2009). An example of the ISR trial structure can be seen in Figure 1, panel B. Stimuli and Psychopy script here: https://osf.io/s68q9/.

### fMRI data acquisition

Structural and functional MRI data were collected at the York Neuroimaging Centre, University of York, using a 3T GE HDx Excite MRI scanner with an eight-channel phased array head coil (GE) tuned to 127.4 MHz, across two scanning sessions run on different days. We acquired a structural MRI for each participant based on a T1-weighted 3D fast spoiled gradient echo sequence (TR = 7.8 ms, TE = minimum full, flip angle 20°, matrix size = 256 x 256, 176 slices, voxel size = 1.13 x 1.13 x 1 mm^3^). Task-based activity was recorded from the whole brain using single-shot 2D gradient-echo echo planar imaging (EPI) from 38 axial slices aligned with the temporal lobe (flip angle = 90°, matrix size = 64 x 64, and field of view (FOV) = 192 x 192 mm^2^, TR = 2000ms, TE = Minimum full, voxel size = 3 x 3 x 3 mm^3^, interleaved acquisition). The phonological, semantic and case judgement data was collected across two scanner runs in one session (11 minutes, 330 volumes; following a resting state scan). The ISR task was collected across three scanner runs (11 minutes, 330 volumes). This was followed by a brief semantic localiser task (used to identify target sites for TMS in the left ATL), which presented words forming sentences or nonwords, one item at a time for passive reading (details in Vatansever et al., 2017). This data was collected in a single scanner run (4 minutes, 120 volumes). An intermediary FLAIR scan with the same orientation as the functional scans was collected in each session between experimental runs to improve the co-registration between subject-specific structural and functional scans.

### fMRI data pre-processing and analysis

fMRI analyses for all fMRI datasets were conducted at the first and higher level using FSL-FEAT version 5.0.8 (Woolrich et al., 2001; Smith et al., 2004; Jenkinson et al., 2012). Pre-processing included slice timing correction, linear motion correction (Jenkinson et al., 2002), high-pass temporal filtering (sigma = 100s), brain extraction (Smith, 2002), linear co-registration to MNI152 standard space (Jenkinson and Smith, 2001), full-width-half-maximum (FWHM) spatial smoothing using a Gaussian kernel of 5mm and grand-mean intensity normalisation of the 4D dataset by a single multiplicative factor.

#### First-level fMRI analyses of phonological, semantic and case judgement tasks

The two scanner runs were modelled independently at the lower level and combined using fixed effects higher-level analyses (FLAME). The pre-processed time series data were modelled using a general linear model correcting for local autocorrelation (Woolrich et al., 2001). Each experimental condition (phonological, semantic and case decision) was modelled using event-based explanatory variables (EV) as a boxcar function, convolved with a hemodynamic response gamma function, in a variable epoch design (Grinband et al., 2008): the start of each epoch was defined as the onset of the condition stimulus, with epoch duration determined by the response time on each trial. This approach was selected to control for time-on-task effects since response times differed between localiser conditions and groups. Only correct trials were modelled. Seven contrasts were defined: individual conditions > rest (phonological, semantic and case decisions), phonological and semantic decisions > case decisions and the contrast between the phonological and semantic tasks (which was used to select stimulation sites in the TMS experiment).

#### Second-level fMRI analyses of phonological, semantic and case judgement tasks

After combining runs in a fixed effects design, we ran two sets of second level analyses using a mixed effects design whole-brain analysis (FLAME http://www.fmrib.ox.ac.uk/fsl): First, an omnibus GLM modelling each judgment condition and contrast at the overall group-level was run to determine the whole-brain main effects of the judgment tasks across all participants. Second, since we expected patterns of functional recruitment related to semantic and phonological judgments to vary between our participants based on their behavioural profile, individual differences in whole-brain activation related to the continuous semantic reliance measure were analysed in separate regressions for each judgment task with the z-scored semantic reliance predictor (detailed earlier). All analyses were cluster corrected using a z-statistic threshold of 2.6 to define continuous clusters. A pre-threshold grey-matter mask was applied to constrain analyses to cortical and subcortical grey matter. Multiple comparisons were controlled using Gaussian Random Field Theory (RFT) at a significance threshold of p <.05.

#### First-level fMRI analyses of immediate serial recall

The three runs were modelled independently at the lower level using a general linear model correcting for local autocorrelation (Woolrich et al., 2001). In order to isolate rehearsal-related activity, activation was estimated at each voxel using regressors modelling the encoding, rehearsal and retrieval phases for each condition separately (for the correct nonword, random word, semantic sequence and visual baseline trials). Thirteen rehearsal-epoch contrasts were defined: individual conditions (rehearsal of nonwords, random words, semantic sequences and visual baseline stimuli) over rest; nonwords, random words and semantic sequences over the visual baseline; plus contrasts of nonword over word rehearsal, nonword over semantic sequence rehearsal and word over semantic sequence rehearsal (and the reverse contrasts).

#### Second-level fMRI analyses of immediate serial recall

The three runs were combined using a fixed effects design, and second level analyses were performed using mixed-effects whole-brain analysis (FLAME). Like the judgment tasks, we ran an omnibus GLM to determine whole-brain correlates of experimental manipulations across all participants (i.e., modelling the encoding and rehearsal epochs of each immediate serial recall condition and the contrasts of interests across the group). We also examined the whole-brain correlates of individual differences in semantic reliance in separate GLMs for each ISR rehearsal manipulation (i.e., separately modelling each rehearsal condition over rest and including individual z-scored semantic reliance as a predictor). All statistical thresholds were identical to analyses of the word judgement tasks.

#### Region-of-interest analyses testing the functional specificity of semantic reliance effects

In order to visualise how significant activation related to the continuous regressor broke down at the SR group level, we conducted region of interest analyses of clusters identified at the whole-brain level for any experimental conditions (judgement or rehearsal conditions over rest) associated with our behavioural semantic reliance regressor. In each case, we used the FEATquery tool in FSL to extract each individual’s unthresholded percentage signal change within a binarised mask of the significant semantic reliance-modulated cluster for each experimental condition. The extracted data for these clusters were used to assess correlated behavioral and between-task activity. Given the modest sample size, Pearson correlations were accompanied by bootstrap 95% confidence intervals (5000 resamples; bias corrected) to provide inference robust to non-normality and outliers.

#### Psychophysiological interaction (PPI)

Following identification of a cluster within left anterior inferior parietal cortex overlapping with left dorsal supramarginal gyrus (SMG) in analyses of the word judgement tasks that was related to individual differences in semantic reliance, we extracted the time-course from a binarised mask of the group-level cluster, transformed into each individual’s native space, to look for psychophysiological interactions (PPI; O’Reilly et al., 2012). These effects reflect differences in functional coupling of anterior inferior parietal cortex with other brain areas according to the task condition and semantic reliance of participants. For both the word judgement and immediate serial recall data, the extracted time-courses of left anterior inferior parietal cortex and the interaction term were included in GLM models as explanatory variables (at the lower level, for each participant and each run individually). These were then submitted to group-level analyses (examining the contribution of semantic reliance for each word judgement condition and each immediate serial recall condition), using the same contrasts and thresholds as the whole-brain analyses above.

#### Intrinsic connectivity analysis from resting-state fMRI (independent data set)

Resting-state functional connectivity analyses were used to establish patterns of intrinsic connectivity for the anterior inferior parietal cortex site that showed effects of semantic reliance. This analysis was done in an independent sample of resting-state fMRI in 195 healthy students from the University of York; a dataset that has been used in multiple previous publications (e.g., Sormaz et al., 2018; Vatansever et al., 2017a; Zhang et al., 2021). We removed participants from this sample who were within our low-and high-semantic reliance sample.

Resting-state functional connectivity analyses were conducted using CONN-fMRI functional connectivity toolbox, version 18a (http://www.nitrc.org/projects/conn) (Whitfield-Gabrieli and Nieto-Castanon, 2012), based on Statistical Parametric Mapping 12 (http://www.fil.ion.ucl.ac.uk/spm/) and followed the processing steps detailed in Zhang et al. (2021). The resulting nii.gz maps were overlaid with the SR-moderated PPI data to examine overlap.

### TMS experiment

The TMS experiment was designed to investigate the effects of transient SMG disruption on short-term recall performance in participants with high and low semantic reliance. Given that activation in anterior inferior parietal cortex overlapping with SMG was associated with semantic reliance in the task-based fMRI studies, we reasoned that immediate serial recall might be differentially disrupted by inhibitory TMS applied to SMG depending on the semantic reliance of the participants and the semantic and phonological demands of the task. In a separate session, the effect of TMS to a second site (left lateral ATL; anterior portion of MTG) was also tested. Since left ATL is associated with semantic but not phonological processing, inhibitory stimulation of left SMG and left ATL should elicit dissociable effects.

A subset of participants from the fMRI studies completed the TMS sessions. They were tested on their auditory-verbal recall of lists of words and nonwords and a control visual pattern memory task before and after 40 seconds of continuous theta burst TMS applied to an individually localised site within left SMG. These behavioural tasks were designed to be short, lasting for around 10 minutes in total, to ensure that any effects of TMS would be strong during the period of data collection. To optimise experimental sensitivity to effects of TMS, task difficulty was established on an individual basis by presenting lists for recall that were one item beyond individual recall span capacity (for word, nonword and visual pattern span tasks; following Savill et al., 2019a).

#### Immediate serial recall stimuli for TMS study

Forty lists of words and forty lists of nonwords for immediate serial recall were taken from Savill et al. (2019a). We presented twenty lists of each type before and after TMS. A further four lists were created for practice trials before the experiment. Since we adjusted the length of these lists according to participants’ performance, eight-item lists were generated to accommodate individuals with high spans. Lists comprised unrelated monosyllabic items, with a consonant-vowel-consonant structure, and were constructed such that no phoneme was repeated within a list in each syllable position. Word lists comprised unrelated nouns and were allocated to two sets (presented pre- and post-TMS) that were matched for lexical frequency and imageability (Table 1). Nonword lists were created by recombining the phonemes of the resulting word lists (e.g., the word list “bus, note, patch, hawk, yell, roof, game, dish” generated a nonword list putch, hoce, yal, norb, ket, roosh, gim, dafe, or /p□tʃ/ /hə□s/ /jæl/ /nɔb/ /ket/ /ruʃ/ /g□m/ /de□f/ in the international phonetic alphabet). Stimuli were individually recorded by a female British English speaker and edited to 750ms in length, with background noise removed and average intensity controlled using Praat (www.praat.org).

**Table 1.** Average properties of the two sets of word lists used within a TMS session.

|  | List Set A | List Set B |  |
| --- | --- | --- | --- |
|  | *M (SD)* | *M (SD)* |  |
| Lexical Frequency | 4.13 (0.22) | 4.09 (0.23) | $t(38) = 0.47, p = .63$ |
| Imageability | 5.67 (0.33) | 5.69 (0.29) | $t(38) = -0.24, p = .81$ |

*Note.* List Set A and B were allocated to TMS baseline and post-TMS conditions in a counterbalanced fashion, between participants. Stimuli were reordered within their lists for the second TMS session tested at least 7 days later. Lexical frequency values = SUBTLEX Zipf Frequencies (van Heuven et al., 2014); Imageability values = ratings from Cortese and Fugett (2004).

#### Visual pattern memory task stimuli for TMS study

The effects of TMS on immediate serial recall were compared with a control short-term memory task that did not involve language processing. Participants were asked to remember visual patterns comprising grids of cells; approximately half of the cells were shaded (cells ~ 4 × 3 cm on screen), and participants were asked to remember this visual pattern (see Figure 3). As for the immediate serial recall tasks (and the procedure for the equivalent TMS task in Savill et al., 2019a), we identified the array size that participants could remember correctly, and then presented arrays in the TMS study that were one grid size larger than this capacity, to maximise task sensitivity for individuals. Two non-contiguous black cells in each pattern array were changed to white to create test trials: participants were asked to identify the locations that had changed. Forty unique pattern arrays were presented in the TMS sessions (10 pre- and post-TMS for the SMG and ATL stimulation sessions).

**Figure 3.**
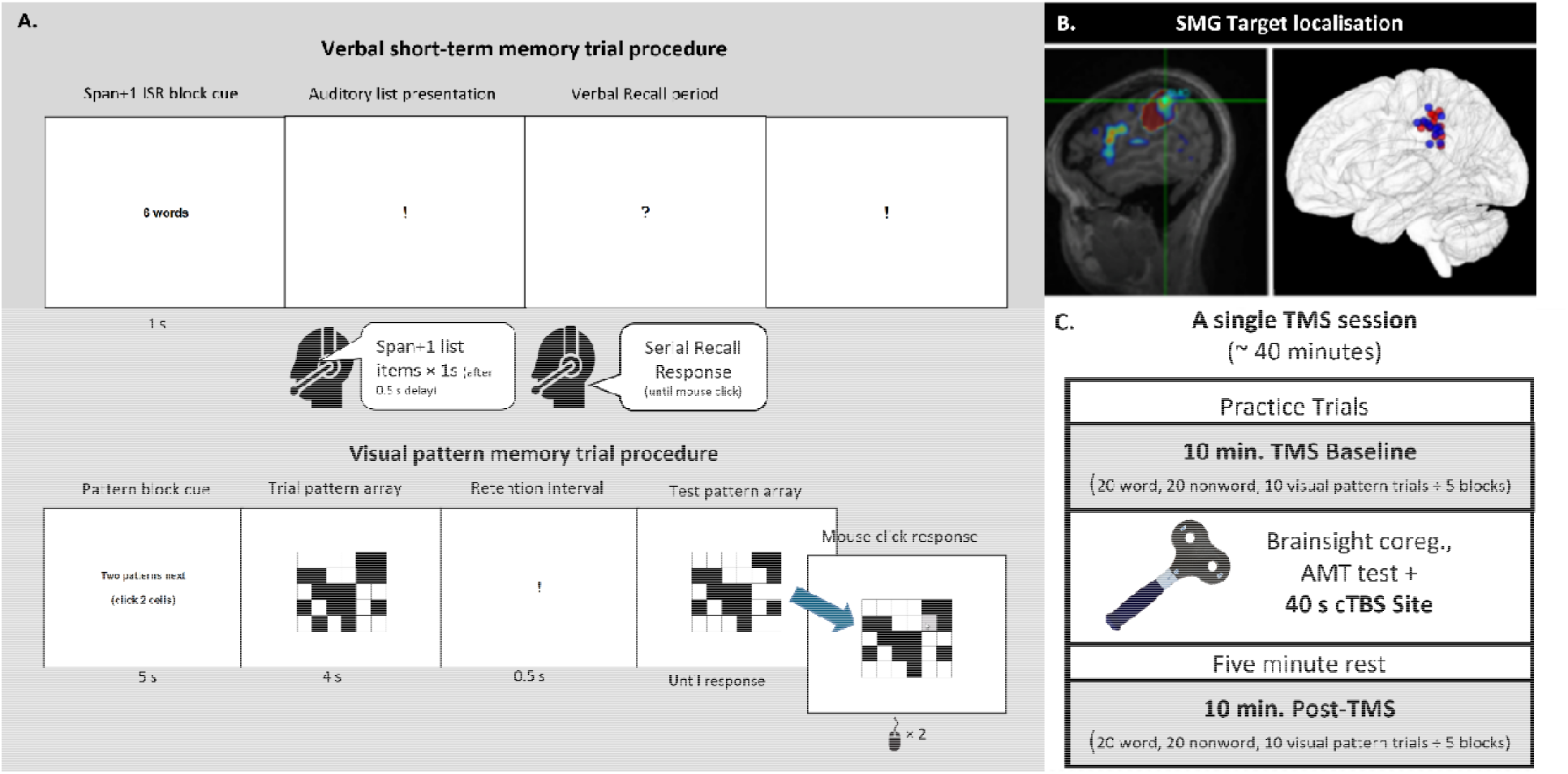
TMS Experimental Protocol. (A) The structure of immediate serial recall and visual pattern memory trials. The visual display is shown above the concurrent auditory components of the verbal short-term memory tasks. The procedure for word and nonword trials differed only in the number of items presented in each list. Participants were instructed to try to verbally recall all the items in order at the end of each list, identified by a question mark on screen. In visual pattern memory trials, participants clicked the two white cells in the test pattern array that were black in the trial pattern array. (B). Peak fMRI activation was used to identify TMS targets in regional masks transformed to individual space. The left panel shows an example of a participant’s phonologically-localised SMG site, within the SMG mask shown in red. The right panel shows the target sites across participants (red spheres = high semantic reliance; blue spheres = low semantic reliance). (C). Structure of a single TMS session. Different experimental sets were used to test performance in the baseline and post-TMS phases.

#### TMS span testing

Prior to the first TMS session, participants’ word and nonword spans and their visual pattern memory capacity were assessed. Word and nonword lists of increasing length (four lists per length, from three to eight items) were auditorily presented and participants were asked to repeat back each list in order. Span sizes for words and nonwords were determined as the final length that at least two of four lists were recalled completely correctly. Word and nonword lists containing one item above the span length of each participant were presented in the TMS sessions. Visual pattern memory capacity was established in a similar way: four trials of each grid size were tested, gradually increasing in size (3×3, 3×4, 4×4, 4×5, 5×5, 5×6, 6×6, 6×7, 7×7, plus additional 5×5, 5×6, 6×6, 6×7 pattern arrays each with one fewer black cell). The grid size selected for the TMS experiment was one size larger than the final grid size at which at least two of four trials had been fully correct for each participant. The final list lengths presented in the TMS sessions ranged from 5-7 words and 4-5 nonwords. The final visual pattern arrays ranged from 5×5 to 6×7 cells. Test sizes did not differ between SR groups (details in Supplementary Materials).

#### Localisation of TMS stimulation sites

Stimulation sites were identified using the Brainsight TMS-MRI co-registration system (Rogue Research, Montreal, Canada, www.rogue-research.com). Peak responses within individuals’ fMRI data were selected within anatomical masks corresponding to left SMG and ATL (using the Harvard-Oxford atlas in FSL), transformed into native brain space. Individual unthresholded functional maps in individual space were overlaid on participants’ T1 structural images (example in Figure 3B). The target site for SMG stimulation was identified by selecting the peak phonological judgement activation (using the contrast phonological judgement > baseline) within the transformed SMG mask. Individual SMG target sites for the group are plotted in Figure 3B (average MNI coordinate of SMG targets in standard space was −55, −35, 38). A similar procedure was followed to identify a semantic target site within the left ATL; here we used the sentence > nonwords (*meaningful* > *meaningless*) contrast from the passive semantic localiser task (cf. Vatansever et al., 2017a; further details are provided in the Supplemental Materials Section D). The localised coil targets were marked on a tight-fitting elastic cap worn by the participant throughout stimulation.

#### TMS experimental procedures

Participants completed one version of each task before TMS (to provide a behavioural baseline) and a second version starting five minutes after TMS was delivered. The tasks were run in E-Prime 2.0 (Psychology Software Tools, 2012), using a script from Savill et al. (2019a). Practice trials (two of each type; set to the participant’s test size and containing different stimuli) were used to familiarise participants with the task at the beginning of each session. Participants were told the length of the word and nonword lists and the structure of the testing session: four lists of words were followed by four lists of nonwords (or the reverse, counterbalanced across participants) and then two visual pattern memory trials. These tasks were cycled for approximately ten minutes. Participants wore a headset with an integrated microphone to record spoken responses. For the immediate serial recall tasks, they were asked to attempt to repeat aloud all spoken items, in order of presentation, immediately at the end of each list, and to attempt to recall items even if unsure. For the visual pattern trials, they were asked to identify the two cells in each trial that changed from black (in the target array) to white (in the test array).

A task cue screen appeared at the beginning of a block (i.e., before each set of word lists, nonword lists and visual pattern memory trials; see figure 3). The cue screen displayed for one second for word and nonword lists, and five seconds for the cue for pattern trials to accommodate adjustment to the switch in input modality and task demands. Within a block, a 0.5s preparation screen (exclamation mark fixation) preceded the onset of each trial stimulus. Auditory stimuli were presented at a rate of one item per second, accompanied by an exclamation mark on the screen, which changed to a question mark after the final item in the list had been presented. After recalling each list, participants clicked a mouse button to start the next trial. Responses were digitally recorded for later transcription and coding. In pattern trials, grids of black and white cells were visually presented for four seconds followed by an exclamation mark fixation for 0.5s seconds and then a test array. Participants clicked the two cells they thought had changed to white using the mouse; these cells briefly changed colour to grey to acknowledge each click. The accuracy of clicks was automatically recorded. The test array remained on screen until the second response, and the next trial began.

#### TMS protocol

We used a continuous theta burst stimulation (cTBS) protocol, since this approach has been shown to elicit long-lasting behavioural suppression in previous studies (Huang et al., 2005; Wischnewski and Schutter, 2015). The stimulation lasted for 40s and consisted of bursts of 3 TMS pulses delivered at 50Hz. Bursts were delivered every 200ms (5Hz) for a total of 600 pulses (Huang et al., 2005). Stimulation was administered with a 50mm figure-eight stimulation coil connected to a Magstim Rapid2 stimulator (The Magstim Company, Carmarthenshire, Wales, UK) in a temperature-controlled room. Pulse intensity was set at 80% of individuals’ active motor threshold (AMT), which was assessed immediately prior to cTBS. AMT was determined as the lowest stimulation intensity required to elicit visible contraction of the first dorsal interosseous muscle in the contralateral hand. In participants with an AMT greater than 64% of stimulator output (six participants in each group) the intensity was set at the maximum allowed by the stimulator for this protocol (51%). Stimulation intensities were comparable between sessions (SMG *M* = 48% stimulator output; *SD* = 5.21; ATL *M* = 48% stimulator output; *SD* = 5.04). After cTBS, participants rested for five minutes before starting the recall task to allow the cTBS effect to reach its maximum level (Huang et al., 2005).

#### Verbal recall transcriptions

Recordings of verbal responses were transcribed phoneme by phoneme by an experienced researcher who did not know which recall attempts were pre- and post-TMS. The transcription procedure and coding of errors was identical to that reported in our previous studies (Savill et al., 2015, 2019b).

#### Analysis of TMS effects on immediate serial recall

ISR response analyses focused on the probability of successful item recall on each trial, where target items could be recalled in any list position (cf. Savill et al., 2015). Pattern recall analyses focused on the probability of correctly identified pattern changes for each trial of the task (i.e., correct cell clicks). ISR data and pattern recall data were analysed separately.

As an initial data screen, for each ISR list in each participant, the percentage of target items recalled from the list (adjusted to the length of the test list) was calculated. Next, ISR data were screened to identify whether any participant’s performance significantly differed between their baseline conditions or showed signs of blanket facilitation (improvement in all tasks and conditions). One participant was excluded on this basis, leaving 11 participants in the low-SR TMS group.

#### Statistical Analysis, ISR data

To control for random effects related to fluctuations in the psycholinguistic properties of lists between trials and possible item-related variation in likely recall success of individual lists – in addition to possible differences in recall capacity not sufficiently accommodated for by our adjustments to span size – we used linear mixed effects modelling. To do this, we used PROC GLIMMIX in SAS (v9.4) to model the probability of successful trial outcomes by specifying a binary distribution function with a logit link function. Fixed effects included TMS stimulation site (ATL and SMG), TMS stimulation (pre- and post-TMS), semantic reliance (high and low group), and stimulus type (word and nonword). For dummy coding of these fixed effects, SMG, post-stimulation, low semantic reliance, word stimuli, and list position, respectively, were treated as the reference levels. We included separate random effects for the intercepts for both participants and list items. Finally, we also included list size and item location in a list as covariates.

#### Statistical Analysis, visual pattern memory data

As with the ISR TMS data analysis, we used PROC GLIMMIX in SAS (v9.4) to model the probability of a completely correct trial response in the pattern task, by specifying a binary distribution function with a logit link function. Fixed effects included TMS stimulation site (ATL and SMG), TMS stimulation (pre- and post-TMS), and semantic reliance (high and low). For dummy coding the fixed effects, SMG, pre-stimulation, and low semantic reliance, respectively, were treated as the reference levels. We included a random effect for the intercept for participants, and we also included grid array size and block order as covariates.

## Results

We first examined activation elicited by phonological and semantic judgements in individuals whose immediate serial recall outside the scanner indicated they were either strongly or minimally reliant on semantic information (with high semantic reliance associated with strong effects of imageability and weak nonword performance in ISR). These analyses identified a single cluster of differential activation in anterior inferior parietal cortex overlapping with left supramarginal gyrus (SMG), specifically the dorsal region of SMG bordering superior parietal lobe, during phonological decisions. Next, we examined the response in this region in a verbal short-term memory task as a function of semantic reliance and the lexical-semantic content of the material held in mind. Third, we used psychophysiological interaction (PPI) analysis to explore how functional coupling of this anterior inferior parietal region overlapping with SMG varied across participants and experimental conditions. Finally, we examined whether inhibitory TMS to phonologically localised peaks within left SMG disrupted immediate serial recall performance in a way that reflected semantic reliance and the lexical-semantic properties of the stimuli.

### In-scanner behavioural data

A table summarising the behavioural data for both localiser and ISR tasks is provided in the supplemental materials (Table S2). Bonferroni corrections were applied in behavioural analyses to correct for multiple pairwise comparisons within a model.

#### Phonological, semantic and case judgements

Participants made few errors overall (9% of all trials; 10% excluding the case judgment control). Responses were slower and less accurate for phonological than semantic decisions [main effect of condition on RT: *F*(1,28) = 27.40, *p* < .001, ηp^2^ = .50; and accuracy: *F*(1,28) = 7.77, *p* = .01, ηp^2^ = .22], despite these conditions being behaviourally matched in pilot testing. Since half of the sample were selected to be highly semantically reliant (and phonologically weak, compared with unselected participants), we observed a main effect of semantic reliance (SR group) on judgement accuracy [*F*(1,28) = 5.06, *p* = .03, ηp^2^ = .15] (but not RT [F(1, 28) = 0.09, p = .78, ηp^2^ < .01]) and an interaction of SR group with condition in both accuracy [*F*(1,28) = 6.34, *p* = .02, ηp^2^ = .50] and RT [ *F*(1,28) = 6.23, *p* = .019, ηp^2^ = .18], driven by relatively poor phonological performance (both less accurate and slower) compared to semantic judgments in the high SR group, while accuracy did not differ between localiser conditions in the low SR group. Case decision performance was similar between groups suggesting the groups were similarly attentive to the task (Low SR accuracy M = 95%, SD = 0.93; High SR accuracy M = 95%, SD = 1.44; Low SR RT M = 0.55s, SD = 0.06; High SR RT M = 0.57s; SD = 0.07).

#### ISR Task

There were few errors overall (8% of trials; 10% excluding the low-level baseline). Errors were more likely in the recall of nonwords and random words, relative to semantic sequences [main effect of condition on Accuracy: *F*(2,56) = 30.11, *p* < .001, ηp^2^ = .52; Bonferroni-corrected pairwise comparisons: nonwords vs. random words: *t*(29) = 0.59, *ns*; random words vs. semantic sequences, t(29) = −9.77, p < .001]. There were also differences between conditions in response times [main effect of condition on RT: *F*(2,56) = 47.70, *p* < .001, ηp^2^ = .63]: Bonferroni-corrected pairwise comparisons showed nonwords were slower than random words: *t*(29) = 5.64, *p* < .001; and random words were slower than semantic sequences: t(29) = 4.68, *p* < .001]. Recall probe accuracy tended to be higher overall in the low SR group than the high SR group [*F*(1,28) = 4.13, *p* = .052, ηp^2^ = .13] but there was no interaction between group and condition for accuracy or RT.

### fMRI analysis of phonological and semantic judgements

#### Whole-brain analyses

Phonological and semantic decisions showed common activation, relative to the implicit baseline, in left inferior frontal gyrus extending into precentral gyrus, orbitofrontal cortex and insula. Both tasks also elicited significant activity in bilateral supramarginal gyri into postcentral gyri and superior parietal cortex and lateral occipital cortices extending to temporal fusiform gyri (Figure 4A). The strongest activity was seen in anterior left IFG (pars triangularis) in the semantic condition and in posterior IFG (pars opercularis) in the phonological condition, in line with previous fMRI and TMS studies indicating anterior-posterior specialisation within IFG (fMRI: Demonet et al., 1992; Fiez, 1997; Poldrack et al., 1999; Bokde et al., 2001; TMS: Gough et al., 2005; Hartwigsen et al., 2015; Hartwigsen et al., 2020). IFG activation is prominent in both judgment conditions when semantic and phonological conditions are contrasted with the low-level baseline of case judgements (Supplementary Figure S2). Both tasks also elicited overlapping activity in paracingulate cortex, which may reflect decision-related control demands common to both tasks (Venkatraman et al., 2010).

**Figure 4.**
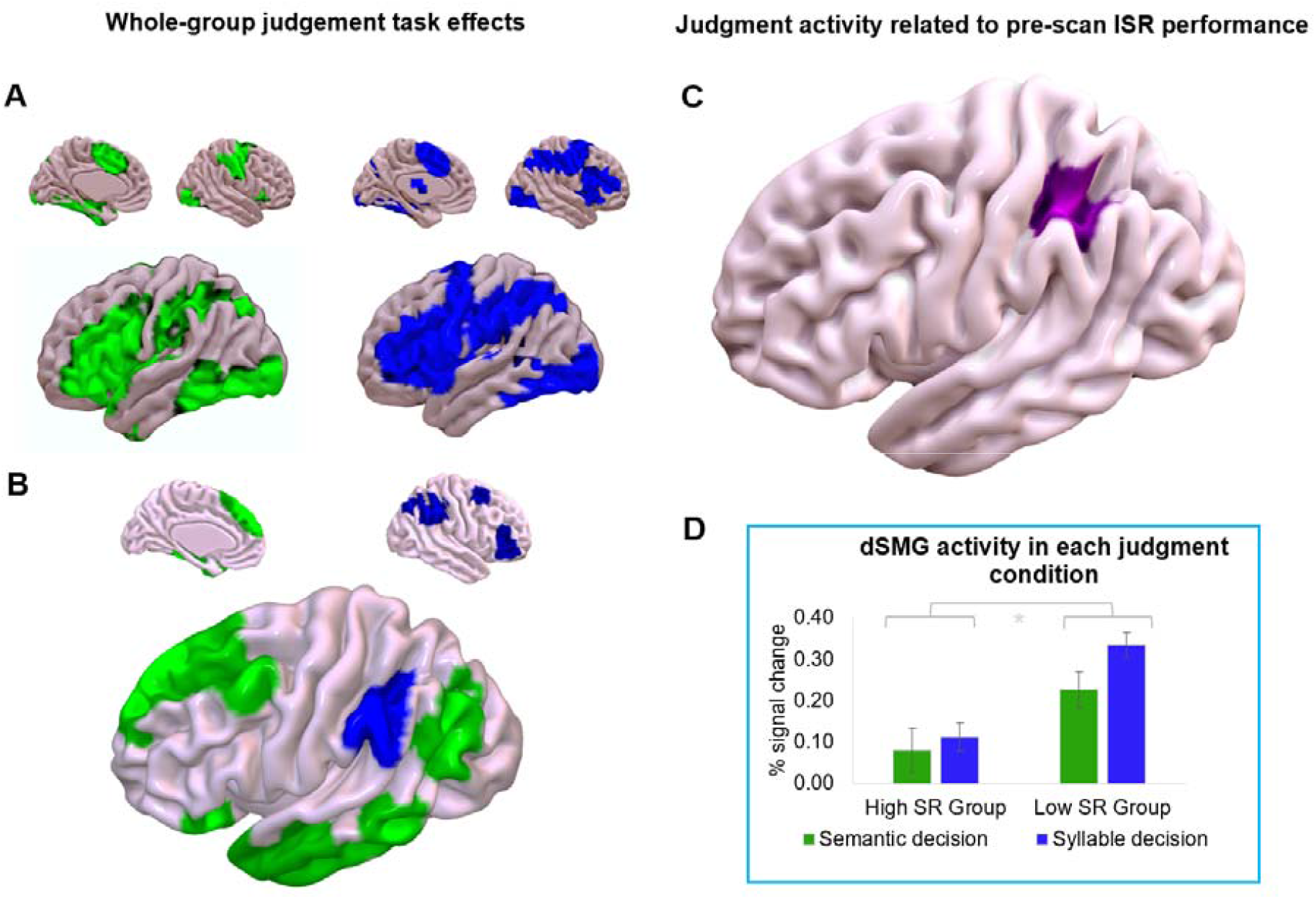
Activation elicited by phonological and semantic decisions. (A) Group level judgment activity over rest. Green: semantic decision > rest. Blue: phonological decision > rest. (B) Clusters corresponding to significant task contrasts (left hemisphere magnified). Green: semantic > phonological decision. Blue = phonological > semantic decision. (C) Cluster in left anterior inferior parietal cortex overlapping with left supramarginal gyrus (SMG), specifically the dorsal region of SMG bordering superior parietal lobe (dSMG), showing activation to phonological decisions that was negatively correlated with behavioural semantic reliance (SR). (D) Percent signal change extracted separately for the two groups of participants (high and low SR) during semantic and phonological decisions from the SMG cluster identified in whole-brain analysis. Participants who were less reliant on meaning to support immediate serial recall showed greater activation of left SMG, especially for the phonological task.

In direct contrasts of semantic and phonological decisions (Figure 4B), the semantic task elicited more activity along the left temporal fusiform cortex, extending into temporal pole and ventral IFG, and laterally into posterior middle temporal gyrus. Semantic decisions also elicited greater activity in left lateral occipital cortex, plus frontal pole, superior and middle frontal gyrus. Phonological decisions, on the other hand, elicited stronger activity in bilateral SMG, with activity in right SMG extending into angular gyrus and lateral occipital cortex. Phonological decisions were also associated with greater activation in right frontal pole and middle frontal gyrus, in addition to activity within left cerebellum. This analysis confirms that our tasks revealed the expected group-level activation for phonological and semantic processing, including greater activation of left SMG (among other sites) for phonological decisions.

#### Associations with individual differences in semantic reliance

We correlated individual differences in the behavioural SR measure with neuroimaging data for the phonological and semantic decisions (relative to rest). These analyses identified a single cluster for the phonological task in the left SMG, bordering the superior parietal lobe (z max MNI coordinates: −44, −42, 48), for which the Harvard-Oxford Cortical Structural Atlas assigned the highest probability to posterior supramarginal gyrus (Figure 4C). Activity in this region was negatively correlated with semantic reliance (z max = 5.06, *p* = .024; *p* = .046, corrected for multiple comparisons involved in a 2-tailed test): individuals with the strongest semantic reliance in STM showed the weakest phonological activation in this region of the brain. Figure 4D shows activation associated with semantic and syllable decisions split by SR score.

Follow-up partial correlations examined whether the effect of SR on activation in SMG was attributable to variance in each of the component measures used to calculate SR (i.e., nonword recall and imageability effects in immediate serial recall). There were significant correlations between SR and phonological activation (percent signal change) within the SMG cluster, even when partialling out nonword recall capacity [r(30) = −.58, p = .001] or the imageability effect [r(30) = −.66 p < .001], confirming that the composite SR measure best captured this difference between individuals.

### fMRI analysis of verbal short-term memory

Rehearsal activity for meaningless and meaningful material in verbal short-term memory. All rehearsal conditions activated frontal pole extending into middle frontal gyrus and inferior parietal cortex including left supramarginal gyrus extending into superior parietal lobe, when compared to the implicit baseline (Figure 5A; the equivalent whole-group contrasts over the low-level baseline condition involving strings of repeated letters are reported in Supplementary Figure S3). Rehearsal of nonword lists over the implicit baseline elicited more extensive activity in left precentral gyrus. Nonword rehearsal additionally activated left inferior frontal gyrus, left insular cortex, bilateral inferior frontal sulcus and supplementary motor cortex, extending into paracingulate gyrus. Word sequence rehearsal elicited significant activation in left insular cortex, posterior middle temporal gyri, left planum temporale and bilateral putamen relative to the implicit baseline. Both rehearsal of nonwords and semantic sequences additionally extensively recruited bilateral cerebellum and angular gyri extending to lateral occipital cortex.

**Figure 5.**
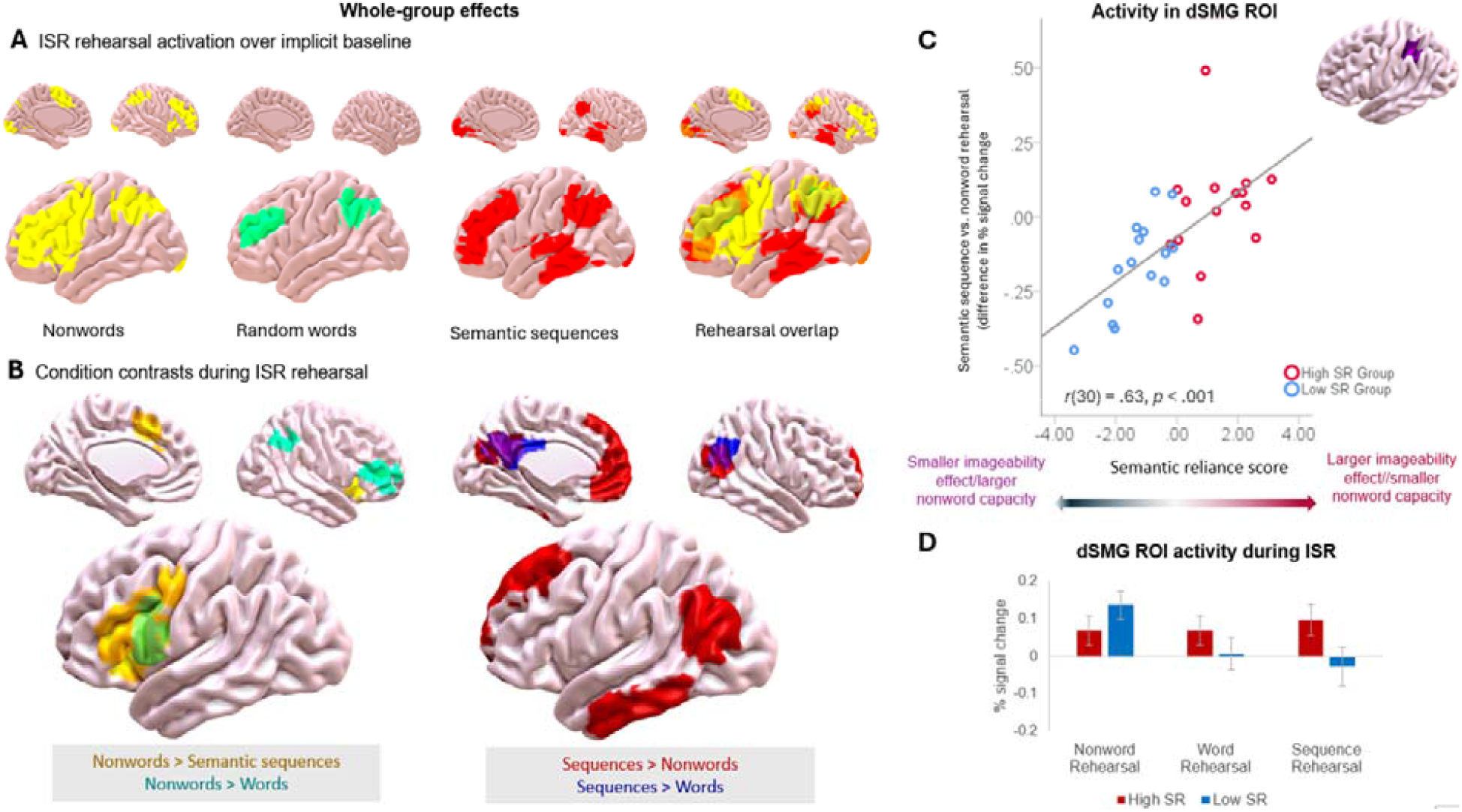
Activation during immediate serial recall (left hemisphere magnified). Panel A shows significant activity over the implicit baseline (yellow = nonword rehearsal; cyan = random word rehearsal; red = semantic sequence rehearsal). Panel B shows significant contrasts between these conditions. Stronger activation for nonwords relative to both word conditions (random words and semantic sequences) is shown on the left (orange-yellow: nonword rehearsal > semantic sequence rehearsal; cyan: nonword > random word; pale green: overlap). Stronger activation for semantic sequences relative to nonwords is shown on the right (red: semantic sequence rehearsal > nonword; blue: semantic sequence > word; purple: overlap). There were no significant clusters reflecting stronger activation for random words compared to the other conditions. C. Difference in signal change between semantic sequence rehearsal activity and nonword rehearsal activity in an ROI within left anterior inferior parietal cortex overlapping with left SMG, focussed on dSMG bordering superior parietal lobe, and defined as the cluster showing an effect of semantic reliance on activation during phonological decisions. Participants who were more semantically reliant showed stronger activation in dSMG for semantic sequences relative to nonwords. D. Activation in each rehearsal condition for participants within the high SR and low SR groups within the left dSMG site defined by examining effects of semantic reliance on activation during phonological decisions. Participants in the high SR group showed similar activation across conditions, while participants in the low SR group only showed activation during nonword rehearsal.

Whole-brain analyses of immediate serial recall identified areas showing activation differences during the rehearsal of nonwords (more reliant on phonology) compared with word conditions (single words and meaningful sequences; Figure 5B). Nonwords elicited stronger activity in left IFG and precentral gyrus compared to both meaningful sequences and random words. Nonword rehearsal also elicited stronger activation relative to random words in right SMG, lateral occipital cortex and right frontal pole, and stronger activation relative to semantic sequences in paracingulate cortex and bilateral insular and orbitofrontal cortices, extending to the right temporal pole. Semantic sequences elicited stronger activation in left anterior inferior temporal gyrus and posterior middle temporal gyrus, bilateral lateral occipital cortex, extending into angular gyrus, and frontal pole, paracingulate and medial frontal cortex. In addition, semantic sequence rehearsal was associated with greater activation in posterior cingulate cortex, precuneus and right lateral occipital cortex compared with the rehearsal of both nonwords and random words (see Figure 5B). The rehearsal of random words did not elicit significantly stronger activity than the other conditions, presumably because neither phonological nor semantic processing was maximised in this condition.

#### Effect of variation in semantic reliance on rehearsal activity

We examined rehearsal-related activity in verbal short-term memory within the left anterior inferior parietal cortex site overlapping with SMG and bordering superior parietal lobe, sensitive to semantic reliance during phonological decision-making (Figure 4C; dSMG). We asked whether the functional contribution of dSMG to verbal short-term memory differed across participants and task conditions. We found that nonword rehearsal activity within dSMG was associated with semantic reliance. Nonword rehearsal elicited stronger dSMG activity than sequence rehearsal [F(1,28) = 4.65, p = .040, ηp2 = .14], and this effect interacted with SR group [F(1,28) = 9.27, p = .005, ηp2 = .25]; greater activation for nonwords relative to semantic sequences in dSMG was specific to the low SR group. Accordingly, there was a strong positive correlation between semantic reliance and differences in rehearsal activity between meaningless and meaningful stimuli in this left SMG ROI [*r*(30) = .63, *p* < 001]: people who were more semantically reliant showed stronger rehearsal activity in this region for meaningful sequences compared with nonwords (see Figures 5C and 5D). Bootstrapped correlational analyses supported the significant negative correlation between behavioural semantic reliance and nonword rehearsal activity in dSMG during ISR in the scanner, r(30) = −.38, p = .04, 95% CI [−.62, −.09]. There was also a significant bootstrapped positive correlation between semantic reliance and differences in dSMG activity between rehearsal of semantic sequences and nonwords, r(30) = .63, p < .001, 95% CI [.38, .82]. This correlation remained even after partialling out the individual contributions of nonword recall capacity [*r*(30) = .40, *p* = .031] and the imageability effect [*r*(30) = .45, *p* = .015], confirming that the composite SR measure best captured differential SMG activity in immediate serial recall.

There were no associations between semantic reliance and SMG activation in other task contrasts. For completeness, we also performed a whole brain analysis to identify any associations between semantic reliance and activation during rehearsal beyond left SMG. This revealed a negative relationship between semantic reliance and nonword rehearsal activity in the anterior putamen: participants who were less reliant on word meaning showed stronger nonword rehearsal activity in bilateral putamen (see Supplementary Materials Section D; Figure S1).

### Psychophysiological Interactions (PPI)

Next, we considered variation in functional connectivity for the left SMG cluster comparing the conditions of the judgement tasks (Figure 6). PPI analyses of the phonological and semantic decisions did not reveal any effects of semantic reliance. Across all participants, however, left SMG connectivity was stronger with precuneus and posterior cingulate cortex in the semantic categorisation condition and this effect extended into right angular gyrus and medial prefrontal cortex when semantic judgements were contrasted with syllable judgements (Figure 6A). These regions revealed by the PPI were largely within the negative intrinsic connectivity map for SMG revealed by resting-state fMRI, indicating that SMG substantially reconfigures its connectivity to support semantic cognition.

**Figure 6.**
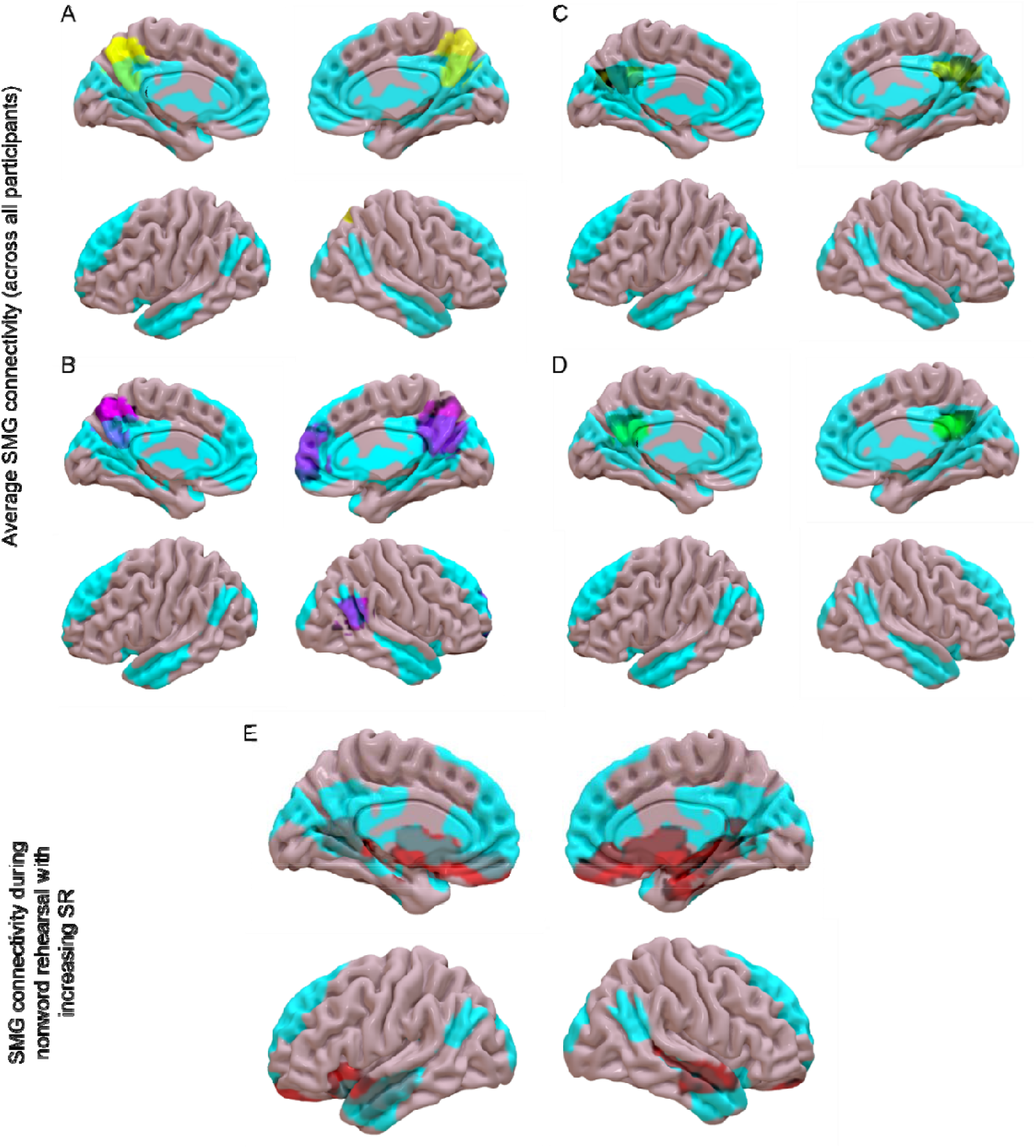
Results of psychophysical interaction models examining how left SMG changes its connectivity across conditions in the judgement tasks (A = semantic judgment > rest, in pale yellow; B = semantic > phonological judgment, in violet) and ISR (C = sequence > nonword rehearsal, in dark yellow; D = word > nonword rehearsal, in green) and how its connectivity varies in nonword rehearsal as a function of semantic reliance (E). All maps are overlaid on the negative intrinsic connectivity of SMG in resting-state fMRI (cyan).

In immediate serial recall, semantic sequence and word rehearsal conditions were associated with greater SMG connectivity in precuneus and posterior cingulate cortex when contrasted with nonword recall across all participants (Figure 6B). This effect overlapped with the connectivity pattern associated with semantic judgements above, showing that SMG to medial parietal connectivity is relevant to semantic processing that supports ISR as well as explicit semantic judgements.

The connectivity of left SMG during nonword rehearsal was also modulated by semantic reliance in medial default mode and left-lateralised semantic regions: stronger semantic reliance was associated with greater connectivity between left SMG and clusters incorporating subcallosal cortex, paracingulate gyrus, orbito- and medial frontal cortex, incorporating left insular cortex, left putamen, bilateral hippocampi and amygdala, lingual gyri extending to left precuneus and posterior cingulate cortex, and bilateral anterior temporal cortex (Figure 6C). These regions were largely within the negative intrinsic connectivity map of SMG, showing that people who are more semantically reliant in ISR also sustain a relatively unusual pattern of SMG connectivity during nonword rehearsal. While the PPI analyses show that semantic judgements, rehearsal of semantic information in ISR and semantic reliance all increase connectivity of SMG to areas it is not normally strongly connected with, the effects of semantic processing across all participants were largely not overlapping with the effects of semantic reliance.

### TMS Experiment

We applied inhibitory TMS to functionally-defined sites in left SMG and left ATL (Supplementary Figure S4) to investigate the causal role of these sites in immediate serial recall as a function of group differences in semantic reliance and the material to be maintained. Our first linear mixed effects model examined the accuracy for each item and included four fixed effects: TMS site (ATL and SMG), TMS stimulation (pre- and post-), semantic reliance (high and low SR groups), and stimulus type (word and nonword), together with interaction terms and covariates (list size and item location). Table S3 (supplementary materials) shows the full set of outcomes from this model; Pearson chi-square by degrees of freedom was .92, suggesting there was no over-dispersion in the fitted model. The 4-way interaction between all fixed effects was significant (*F*(1, 18563) = 4.01, *p* = .045); we explored this interaction through separate analyses for ATL and SMG. These models included three fixed effects, TMS stimulation (pre- and post-), semantic reliance (high and low) and stimulus type (word and nonword), together with interaction terms and covariates (list size and item location). The Pearson chi-square by degrees of freedom for these models were .87 and .88 respectively, indicating no over-dispersion. The key result was a significant 3-way interaction for SMG (*F*(1, 8979) = 5.67, *p* = .02; Table S4), but not ATL (*F*(1, 8973) = 0.19, *p* = .66; Table S5). The modelled recall probabilities (least squares means) for the low and high SR groups are displayed in Figure 7. TMS applied to left SMG affected the immediate serial recall of words and nonwords differently depending on the semantic reliance of the participants. Post-hoc comparisons showed that SMG stimulation disrupted word recall in the high SR group (*t*(8979) = 2.39, *p* = .016), and nonword recall in the low SR group (*t*(8979) = 2.23, *p* = .026), relative to performance in the absence of TMS. Bootstrapped versions of these analyses identified the same patterns and are reported in the Supplementary Materials (Section F).

**Figure 7.**
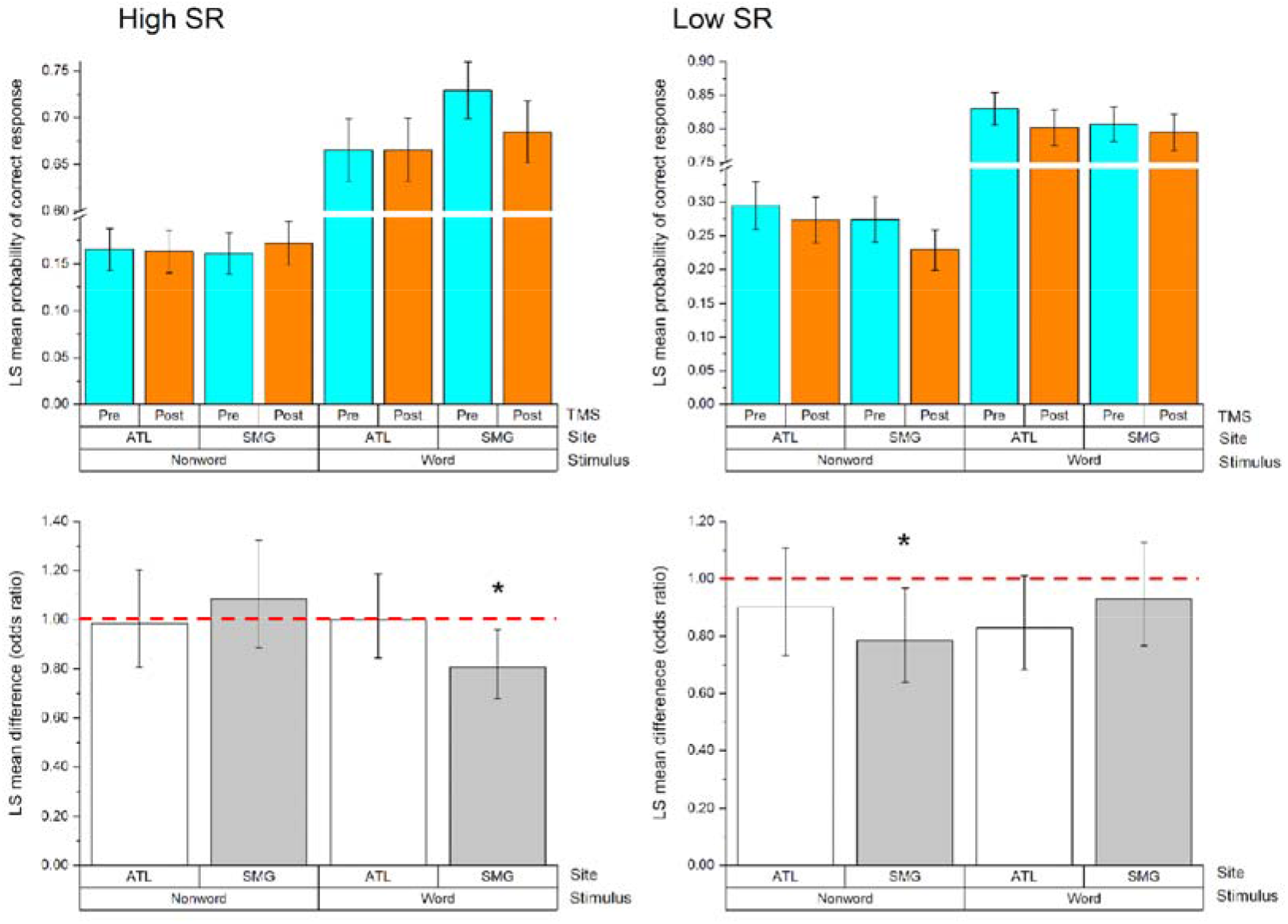
Effects of cTBS TMS. The bar graphs in the top panel show the probability of a correct recall response, before and after TMS, for participants selected to show high semantic reliance (SR) (left side) and low SR (right side). The bottom panel shows predicted mean changes in recall performance following TMS compared to baseline, for nonwords and words, separately for ATL and SMG. These data are expressed as odds ratios, since in a logistic model, the effect of TMS can be characterised in terms of changes in the probability that a particular item or pattern will be recalled. Differences in odds ratios less than one reflect a reduced probability of recall following TMS (1 = no change from pre to post-TMS; red dotted line provided for reference). Error bars represent 95% confidence intervals. * = *p* < .05.

There was no effect of TMS on visual pattern memory (see Table S6 for the linear mixed effects model outcomes), indicating that the disruptive effect of SMG stimulation on immediate serial recall cannot be attributed to general effects of stimulation.

## Discussion

We delineate a key neurobiological marker of individual differences in verbal short-term memory across fMRI and TMS. Behavioural research has shown that, in immediate serial recall, poorer performance for nonwords is associated with stronger effects of imageability when repeating lists of words – participants showing this pattern are thought to be more reliant on word meaning or imagery when maintaining verbal information in mind (Savill et al., 2019b). We scanned participants showing high and low semantic reliance in verbal short-term memory as they performed phonological and semantic judgements. Left anterior inferior parietal cortex, associated with phonological function, showed a stronger response to syllable than semantic judgements as expected; however, the response of this region also varied across individuals according to their semantic reliance (with this metric reflecting higher imageability effects and poorer nonword performance in ISR): people with higher semantic reliance showed attenuated responses during phonological judgements. In a second fMRI experiment, the same participants were asked to hold in mind lists of nonwords and words, which comprised either unconnected words or semantic sequences: the left anterior inferior parietal region implicated in semantic reliance in the first study also showed stronger activation during the maintenance of semantic sequences in people more reliant on meaning. Functional connectivity analyses revealed that individual differences in the recruitment of this region are likely to reflect variation in connectivity: for semantically-reliant participants during nonword rehearsal, this site showed stronger connectivity with default mode network regions implicated in semantic processing, including bilateral ATL and ventromedial prefrontal cortex – even though these regions are generally anti-correlated with left dorsal SMG at rest. Finally, while inhibitory TMS stimulation of SMG disrupted verbal short-term memory for nonwords in people with low semantic reliance, for semantically reliant participants, TMS impaired the immediate serial recall of word sequences, demonstrating that this site makes a necessary contribution to different aspects of short-term memory across individuals. These results show that individual differences in the balance of phonological and semantic processes not only reflect a division of labour between semantic and phonological sites, as envisaged by interactive-activation models of language processing, but also relate to the functional tuning and connectivity of an individual region of cortex.

Regions within inferior parietal cortex (IPC), including the supramarginal gyrus (SMG) and angular gyrus (AG), show graded functional and anatomical organization supporting multiple cognitive domains (Humphreys and Lambon Ralph, 2015; Graves et al., 2023). SMG is classically associated with phonological processing and stimulus-driven attention, while AG is more reliably engaged during semantic and episodic retrieval (Binder and Desai, 2011; Humphreys and Lambon Ralph, 2015). These functional specialisations are mirrored by differences in connectivity: SMG shows stronger links to auditory language areas and motor speech regions, whereas AG is more integrated with default mode network regions implicated in long-term memory and heteromodal conceptual processing (Humphreys and Lambon Ralph, 2015). However, both sites fall within a broader ventral parietal system that buffers information over time, supporting the maintenance and integration of multimodal inputs (Humphreys et al., 2020). Phonological buffering of rapidly evolving inputs is supported by SMG when dealing with unfamiliar, unpredictable inputs such as nonwords, while the semantic buffering of multisensory information over longer-time scales in AG supports the retrieval of long-term knowledge (Humphreys and Lambon Ralph, 2015). This pattern is consistent with our group-level results showing greater SMG activity for phonological decisions and nonword rehearsal, and greater AG activity for semantic tasks.

We also observed increased MDN activation, including in intraparietal sulcus and right PFC, during more demanding nonword maintenance conditions, likely reflecting the contribution of cognitive control when automatic language buffering mechanisms (Oberhuber et al., 2016; Majerus, 2019) are insufficient for task performance (Jefferies et al., 2004). Conversely, semantic conditions elicited more activation in default mode regions, consistent with engagement of long-term memory representations (Binder and Desai, 2011; Wirth et al., 2011; Murphy et al., 2018), although relative task difficulty may also have contributed to these effects (Mckiernan et al., 2003; de Dreu et al., 2019).

Despite these group-level distinctions, our data indicate that functional tuning within the left inferior parietal cortex is not fixed but shaped by individual differences in connectivity and task strategy. In participants with higher semantic reliance, SMG responded more to meaningful stimuli compared to nonwords and adopted a more semantic-like connectivity profile. Similarly, while the behavioural effects of TMS were relatively weak in this group, SMG-TMS disrupted word rather than nonword recall in participants with high semantic reliance (contrary to low-semantic reliance participants and typical findings, e.g. Savill et al. 2019). Rather than reflecting discrete cortical regions specialised for phonological or semantic processing, there appears to be functional overlap between these domains (Mattheiss et al., 2018). While some accounts describe SMG as a relatively specialised phonological buffer, drawing on decoding and lesion evidence (Purcell et al., 2021; Yue and Martin, 2021), others link it to broader roles in cognition, such as short-term buffering within and beyond language (Deschamps et al., 2020). These varying accounts can be reconciled within a framework that sees SMG as a flexible store of dynamically evolving inputs, tuned by transitional structure and shaped by graded connectivity across the parietal surface (Humphreys and Lambon Ralph, 2015; Humphreys et al. 2020). This account accommodates both the group-level specialisation observed across studies and the individual variability we report, without positing sharply defined functional subdivisions. Instead, functional engagement emerges from a continuous interaction between intrinsic connectivity, task context, and cognitive strategy, with SMG well-placed to adaptively support either phonological or semantic processing based on input properties and the internal resources available to the individual.

An important point for consideration regarding interpretation of the adaptive activation in anterior IPC is that stimulus presentation in fMRI was visual, unlike the auditory-verbal format of the short-term memory tasks used to select participants and test with TMS. While this helped to eliminate potential acoustic masking effects of scanner noise, it will have required an initial orthographic-to-phonological translation before maintenance can occur. Importantly, the fMRI ISR paradigm was optimized to isolate activity during the subsequent *rehearsal* phase – our primary focus – rather than stimulus encoding. While visual processing might affect the downstream stability of the phonological trace, our findings specifically highlight how dSMG networks manage the active maintenance of these representations once formed.

A limitation of our study is that the group-level dSMG cluster linked to semantic reliance was not spatially identical to the functionally-localised TMS sites. The SMG semantic reliance site was located more dorsally and slightly posterior to the phonological localiser site, adjacent to intraparietal sulcus (IPS). This anatomical position may be informative, however, because the functionally flexible region may lie at the boundary of the core phonological territory, for example, where more ventral phonological and dorsal attentional systems converge. Our data cannot determine whether this functional flexibility across phonological and semantic domains is a property of the broader anterior inferior parietal cortex (IPC), or is specific to the SMG-border territory identified here. Given the fMRI effect in dSMG is proximal to IPS, the differential vSTM activation may also reflect relative attentional engagement and varying recruitment of left ventral and dorsal attention parietal sites (Majerus et al, 2012). This could reflect individual differences in the balance between phonological processing in the dorsal language stream (Saur et al., 2008) and more controlled dorsal attention mechanisms (Majerus, 2025). In any case, the pattern of individual differences was strikingly consistent across both fMRI and TMS modalities: regions within anterior IPC involving SMG showed greater engagement during semantic processing in participants who behaviourally relied more on semantic information in ISR. These findings suggest that functional variability in this region may reflect differences in individuals’ recruitment of semantic versus phonological resources during language tasks.

Future work should investigate the extent and functional significance of this variability across anterior IPC, where graded differences in connectivity are known to link to multiple cognitive networks— including those involved in semantics, phonology, attention, and executive control (Oberhuber et al., 2016). Given that ISR can be supported by multiple cognitive routes, participants may differ in the degree to which they draw on these resources, depending on both stimulus properties and individual cognitive preferences—paralleling prior accounts of individual variability in reading strategies (Hoffman et al., 2015; Woollams et al., 2016). Finally, regions beyond left anterior IPC might also show individual-level shifts in functional engagement that support flexible language processing.

In summary, our findings show that individuals recruit different neurocognitive resources to support verbal short-term memory. By recruiting separate groups of participants selected to show different levels of semantic reliance (or imagery), we were able to show that these behavioural differences reflect different functional tuning within the left anterior inferior parietal cortex, which has been long associated with short-term memory for language. In contrast to previous perspectives, we found that these individual differences reflected divergent connectivity patterns of one cortical region, as opposed to the differential recruitment of distinct large-scale networks underpinning semantics and phonology. The combination of TMS and fMRI demonstrated that this variation in functional organisation had causal consequences for language behaviour.

## Acknowledgments

The project was supported by a grant from the European Research Council (SEMBIND). We would like to thank James Davey, Glyn Hallam and Charlotte Murphy for their help with experimental scripts for fMRI, and Dominic Arnold for assistance with screening participants for inclusion in the high and low semantic reliance groups.

## Data availability statement

Neuroimaging data at the group-level as unthresholded statistical z maps are available in NeuroVault at https://neurovault.org/collections/FLPKKJZP/. Script for the tasks, including TMS task stimuli and analysis script output, and fMRI task thresholded nifti maps are accessible in the Open Science Framework at https://osf.io/s68q9/. The conditions of our ethical approval do not permit public archiving of the raw and individual level neuroimaging data because participants did not provide sufficient consent. Researchers who wish to access the data should contact the corresponding author, N.S., and data will be released to researchers when this is possible under the terms of the General Data Protection Regulation.

