## Supplemental Materials for "Diverse ways of holding verbal information in mind revealed with functional Magnetic Resonance Imaging and Transcranial Magnetic Stimulation: Individual differences in left anterior parietal cortex"

**for**

**Section A: Behavioural data collected in advance of data collection for the present study**

A summary of behavioural screening data for participants invited to take part in the present study is in Table S1.

Table S1. *Screening measure performance in the high- and low- semantic reliance groups*

| | High SR Group | | Low SR Group | | |
|---|---|---|---|---|---|
| | *M* | *SD* | *M* | *SD* | *p* |
| ISR Screening | | | | | |
| Imageability effect in ISR (LI recall % of HI recall) | 72.94 | 5.71 | 97.42 | 6.9 | <.001 |
| High-Imageability Word Recall (%) | 45.02 | 7.94 | 44.02 | 10.09 | 0.76 |
| Low-Imageability Word Recall (%) | 32.82 | 6.12 | 42.77 | 9.73 | <.001 |
| Nonword recall (%) | 24.89 | 8.78 | 32.86 | 8.75 | 0.02 |
| Semantic Reliance score (inverse z-Imageability effect – z-Nonword recall) | 1.3 | 1.04 | -1.3 | 0.92 | <.001 |
| Age (years) | 20.4 | 2.82 | 20.67 | 4.15 | 0.84 |
| Nonverbal Intelligence | | | | | |
| Matrix Reasoning (WASI age-scaled score) | 57 | 7.06 | 56.64 | 6.82 | 0.89 |

| | | | | | |
|---|---|---|---|---|---|
| Reading Ability | | | | | |
| Word Reading Efficiency (TOWRE Word standard score) | 104.53 | 9.13 | 104.47 | 11.21 | 0.99 |
| Literacy Achievement/ Orthographic knowledge | | | | | |
| Reading Achievement (WRAT standard score) | 105.07 | 9.82 | 114.67 | 12.41 | 0.03 |
| Spelling Achievement (WRAT standard score) | 112.2 | 12.54 | 119.64 | 10.75 | 0.1 |
| Semantic Knowledge | | | | | |
| Vocabulary (WASI age-scaled score) | 65.67 | 7.78 | 68.07 | 5.5 | 0.34 |
| Graded Picture Naming (Total correct) | 19.86 | 2.41 | 19.14 | 3.7 | 0.55 |
| Synonyms (% correct) | 76.53 | 6.7 | 75.73 | 10.24 | 0.8 |
| Phonological skill/ short-term memory span | | | | | |
| Forward Digit Span (WASI correct) | 10.13 | 2.26 | 11.27 | 2.15 | 0.17 |
| Backward Digit Span (WASI correct) | 6.4 | 1.64 | 8.4 | 2.2 | 0.01 |
| Spoonerisms (Total) | 10.07 | 1.28 | 10.67 | 1.5 | 0.25 |

| | | | | | |
|---|---|---|---|---|---|
| Phonological decoding efficiency (TOWRE nonword) | 104 | 10.7 | 112.2 | 7.22 | 0.02 |

***Pilot behavioural data.*** Prior to finalising the methodology of the fMRI experimental tasks, these were piloted in an independent sample.

The sequences and stimuli in the localiser task had been originally piloted on 10 random volunteers (mean age 30.9, 6 females) who did not take part in any experimental sessions, in order to objectively verify whether the semantic and phonological decisions had a comparable level of difficulty when tested in standard experimental conditions outside of the scanner. Thus, any behavioural differences observed in fMRI were likely to be due to the nature of the selected participants' response to the task rather than an inherent reflection of task difficulty. Pilot testing revealed no significant differences between the three conditions for reaction time (seconds) or accuracy (RT: Semantic vs. phonological; M .80 vs. .85, t(10) -1.22, $p$ = .254, Phonological vs. Control; M .85 vs. .81, t(10) 0.68, $p$ = .512, Semantic vs. Control; M .80 vs. .81, t(10) -0.33, $p$ = .747; Accuracy: Semantic vs. phonological; M .94 vs. .91, t(10) 0.99, $p$ = .350, Phonological vs. Control; M .91 vs. .95, t(10) -0.94, $p$ = .373, Semantic vs. Control; M .94 vs. .95, t(10) -0.18, $p$ = .864).

### *Section B: Behavioural Data collected in the present study*

Table S2. *In-scanner task performance*

| | High SR | | Low SR | |
|---|---|---|---|---|
| | *M* | *SD* | *M* | *SD* |
| Localiser Task | | | | |
| Accuracy (% of trials) | | | | |
| Syllable Decision | 82.87 | 13.40 | 91.73 | 2.60 |
| Natural Decision | 92.00 | 3.89 | 92.20 | 3.03 |
| Response time (ms) | | | | |
| Syllable Decision | 888.40 | 182.42 | 817.50 | 105.61 |

| | | | | |
|---|---|---|---|---|
| Natural Decision | 756.40 | 151.74 | 770.80 | 74.50 |
| ISR Task | | | | |
| Probe response accuracy (% of trials) | | | | |
| Nonword Lists | 83.89 | 10.43 | 90.28 | 4.07 |
| Word Lists | 85.28 | 6.84 | 86.94 | 5.19 |
| Sem. Sequence Lists | 95.83 | 4.45 | 96.94 | 3.33 |
| Response time (ms) | | | | |
| Nonword Lists | 1294.70 | 204.72 | 1382.00 | 266.39 |
| Word Lists | 1207.40 | 188.24 | 1222.30 | 216.60 |
| Sem. Sequence Lists | 1089.10 | 181.59 | 1178.40 | 176.61 |

### Section C: The calculation of the behavioural semantic reliance regressor

Participants for this study were identified according to the size of their behavioural imageability effect in immediate serial recall in a previous study (cf group data summarised in section A): Those with a relatively large imageability effect (better recall for high imageability words compared to low) or little to no imageability effect (similar recall between high imageability words compared to low) were invited. Since these participants also tended to vary in their phonological STM capacity indexed by their nonword recall, in order to capture relevant variation related to imageability sensitivity and nonword capacity we used a composite measure from this behavioural data for use as a regressor throughout the study, by calculating the difference between their inverse z-scored imageability effect and z-scored nonword performance in the immediate serial recall. As reported, the semantic reliance effects were found not to be accounted for solely by sensitivity to imageability or nonword capacity.

Following the reported analyses conducted using the above metric, to assess whether our results could be an artifact of our choice of SR measure used, we subsequently conducted a principal components analysis on their percentage imageability effect (low imageability recall as a percentage of high imageability recall; here lower values = larger imageability effect) and percentage nonword recall from the original behavioural screening data (i.e. including data from non-scanned participants) to alternatively capture relevant individual variation in behavioural performance and run the equivalent analyses. The resulting PCA-derived factor score is functionally equivalent to our original SR score in the scanned participants ($r = -0.998$); both metrics capture the same underlying behavioral variance. This PCA factor explained 63% of the behavioural variance (Bartlett's sphericity $p = .013$). Unsurprisingly, using this latent factor score yields equivalent results in relation to fMRI activation (in the opposite direction): a significant positive correlation between behavioural semantic reliance and nonword rehearsal activity in dSMG during ISR in the scanner, $r(30) = .38$, $p = .04$, 95% CI [.086, .63] and a negative correlation between semantic reliance and differences in dSMG activity between rehearsal of semantic sequences and nonwords, $r(30) = -.63$, $p < .001$, 95% CI [-.82, -.39]).

#### Section D: Whole brain effects of semantic reliance in ISR

As referred to in the main text, at the whole brain level, we found a differential rehearsal effect related to SR: a negative relationship between nonword rehearsal activity in the anterior putamen and SR (i.e., relatively increased nonword rehearsal activity in bilateral putamen with lower SR; z max: -24, 10, -2).

Follow-up ROI analyses of rehearsal activity within this identified putamen cluster indicated that the differential putamen activity related to SR was specific to rehearsal in the nonword condition: On average, nonword rehearsal elicited more *negative* putamen activity than sequence rehearsal [$F(1,28)= 11.24$, $p = .002$, $\eta p2 = .29$], but this interacted with SR group [$F(1,28)= 5.16$, $p = .031$, $\eta p2 = .16$]; whereby the significant decrease in nonword rehearsal activity in putamen relative to sequences was specific to the high SR group.

However, this differential putamen activity during nonword rehearsal was primarily a phonological effect, rather than an index of SR per se: Applying partial correlational analyses of the SR effect on the rehearsal activity within the putamen similar to the SMG ROI analyses in the localiser task, we determined that the differential SR activity in putamen was closely and positively tied to the nonword recall capacity measure (i.e., phonological capacity): Unlike the effect of semantic reliance in SMG – and compatible with increased putamen activity relating to successful non-lexical phonological function – the relationship between SR and differences in putamen nonword rehearsal activity did not survive once the contribution of nonword recall capacity had been partialled out [$r(30) = -.32$, $p = .10$; the correlation marginally survived after partialling out the individual contribution of the imageability effect, on the other hand, $r(30) = -.37$, $p = .049$].

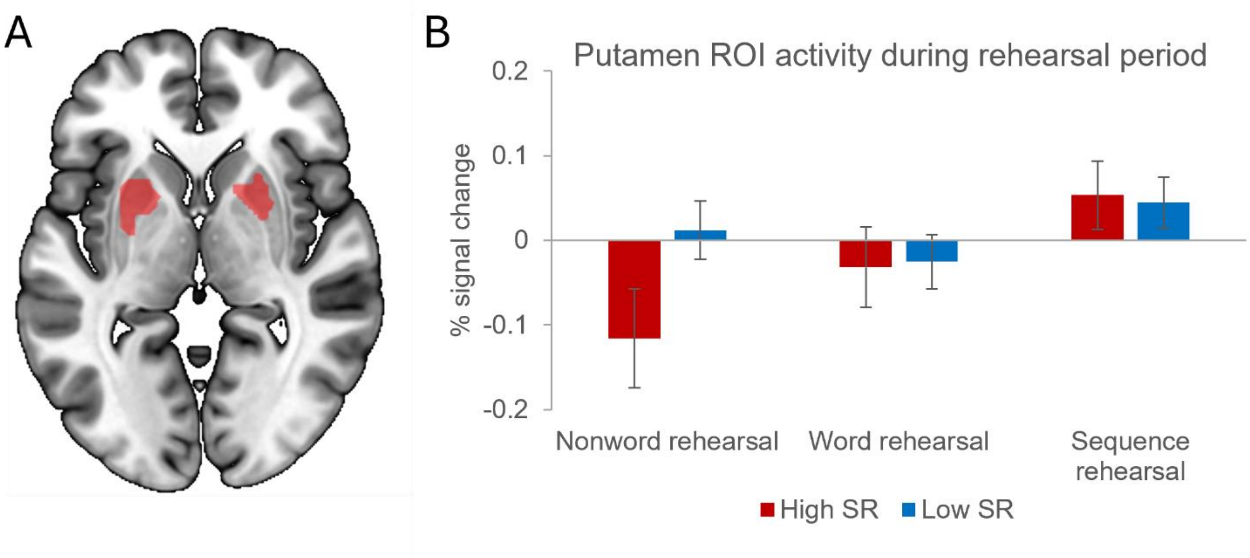


*Figure S1.* Nonword rehearsal activity in the ISR task modulated by the behavioural semantic reliance ISR regressor. Panel A shows clusters of differential nonword rehearsal activity at the whole brain level in anterior putamen correlated with lower SR (red = nonword rehearsal > rest; yellow = nonword rehearsal > semantic sequence rehearsal). B. ROI analysis of activity (signal change from rest/implicit baseline) in the two significant rehearsal conditions, within the SR-defined putamen clusters, according to SR Group.

### Section E: Group task activation contrasts with low-level baseline conditions

In the main text, data in figures 4A and 5A present group task activation over the implicit baseline and subsequent analyses utilise these contrasts. In both the fMRI judgment task and ISR task a low-level control condition was also tested for potential contrast with the experimental conditions. In the judgment task, the baseline condition involved responding whether the unpronounceable letter string was uppercase or not. In the ISR task, the condition denoted 'string' presented four repetitions of the same four uppercase consonants at encoding and at retrieval responding whether the subsequently presented uppercase string corresponded. The results of the corresponding contrasts and maps are provided here.

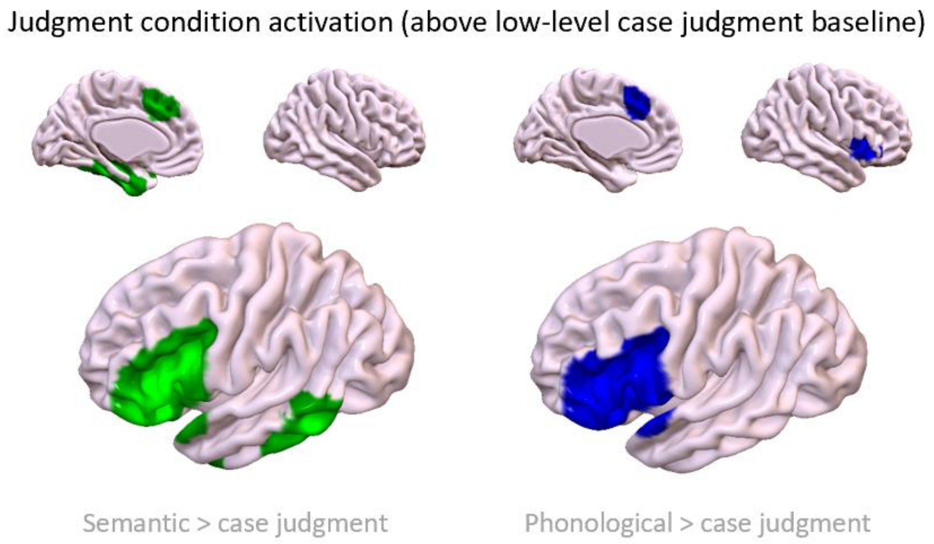


*Figure S2.* Judgement task contrasts with low-level case judgment condition

Activation for both judgment conditions over the low-level baseline was relatively constrained compared to contrasts with the implicit baseline: Activation in both judgment conditions relative to the implemented baseline condition was largely limited to left inferior frontal gyrus, extending to frontal pole and insula; with activity in the semantic condition peaking in anterior left IFG (pars triangularis) and the phonological condition peaking in more posterior IFG (pars opercularis) in line with anterior-posterior specialisation within IFG and linguistic decisions (fMRI: Demonet et al., 1992; Fiez, 1997; Poldrack et al., 1999; Bokde et al., 2001; TMS: Gough et al., 2005; Hartwigsen et al., 2015). The phonological judgment condition also activated right pars opercularis and insula. Both conditions still elicited significant, overlapping activity relative to baseline in paracingulate cortex, which may reflect decision-related control demands common to both tasks compared to the low-level baseline (Venkatraman et al., 2009).

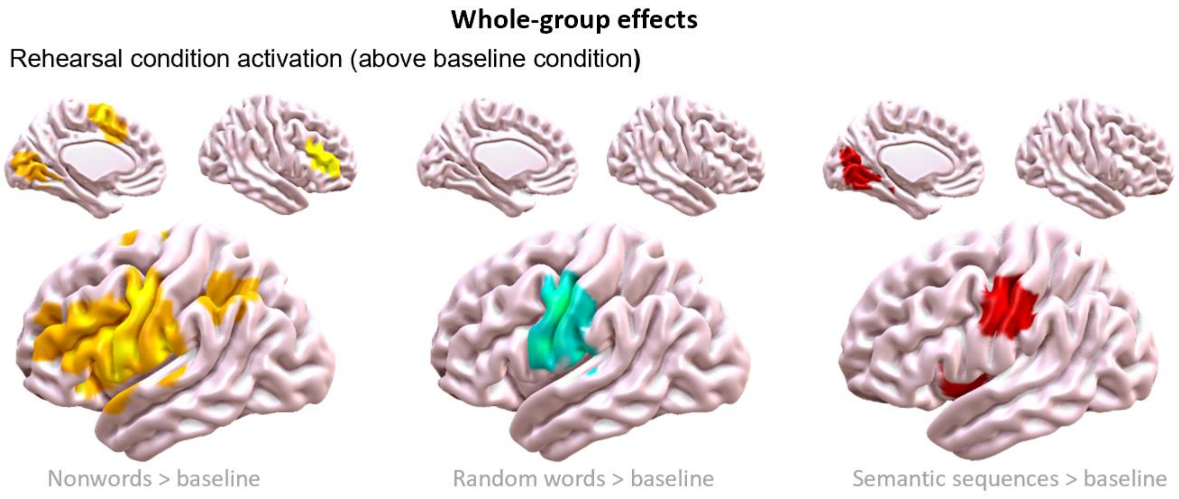

*Figure S3.* ISR rehearsal condition contrasts with low-level string condition

The experimental contrasts of rehearsal conditions over the low-level baseline condition in the fMRI ISR task are shown in Figure S3. All rehearsal conditions activated the cerebellum, the dorsal premotor region of left precentral gyrus, and regions of left postcentral gyrus relative to baseline. In contrast to the baseline, rehearsal of both nonword and random word lists elicited more extensive activity in left precentral gyrus, while both nonword and meaningful sequence rehearsal conditions also activated left insular cortex. Nonword rehearsal additionally activated left inferior frontal gyrus, left posterior supramarginal gyrus, angular gyrus, left paracingulate gyrus and supplementary motor cortex, and frontal pole. Sequence rehearsal elicited significant activation in left putamen relative to baseline. Maintenance of random words did not elicit any other significant activity.

Analyses of whole brain effects of semantic reliance using the respective baseline contrasts yielded no significant activation in either task.

**Section F: Additional information on TMS study**

Prior to participation in the TMS experiment, participants' spans were tested following (Savill et al., 2019) to determine test sizes (next size up from span). Tested word and nonword list lengths and pattern memory grid sizes did not differ, on average, between SR group [Word list length: high SR $M$ = 5.58; $SD$ = 0.67; low SR $M$ = 5.83; $SD$ = 0.58, $t$ (22) = -0.98, $p$ = .34; Nonword list length: high SR $M$ = 4.17, $SD$ = 0.39; low SR $M$ = 4.33, $SD$ = 0.49, $t$ (22) = -0.92, $p$ = .37; Pattern array cells: high SR $M$ = 28.75, $SD$ = 6.94; low SR $M$ = 28.50, $SD$ = 4.21, $t$ (22) = 0.11, $p$ = .92].

The semantic localiser used to identify the ATL stimulation site in the TMS experiment (in place of the categorisation fMRI task) was a passive task first used by Vatansever et al., (2017) (also in Zhang et al., 2019), which has demonstrated reliable activation of brain regions that are expected to be involved in semantic processing (typically from left posterior to anterior medial temporal lobe, and left inferior frontal gyrus). The sentence-reading task used single-shot 2D gradient-echo-planar imaging (TR = 3 s, TE = minimum full, flip angle = 90°, matrix size = 64 × 64, 60 slices, voxel size = 3 mm × 3 mm × 3 mm, 80 volumes). Participants passively viewed meaningful 10 sentences (e.g., her + secrets + were + written + in + her + diary), taken from Rodd, et al https://www.nature.com/articles/s41598-019-52674-9) and meaningless sequences of nonwords (e.g., crark + dof + toin + mesk + int + lisal + glod + flid), item-by-item. The word and nonword sets were matched for both word length and number of syllables and were each presented in two blocks in a pseudo-random order (i.e., a total of 4 blocks). A task instruction (e.g., Meaningful) was used to indicate the transition between different conditions. Each sequence ended with a red fixation lasting 4000–6000 ms. Each word or nonword was presented for 600 ms, followed by a 250 ms fixation before the next item was presented.

Individual semantic site targets for TMS were located using the individual peak activation for the meaningful > meaningless contrast within a lateral temporal mask transformed into native brain space, which largely emerged in superior and middle aspects of the left anterior temporal lobe (stimulation sites for both ATL and SMG shown in Fig S2).

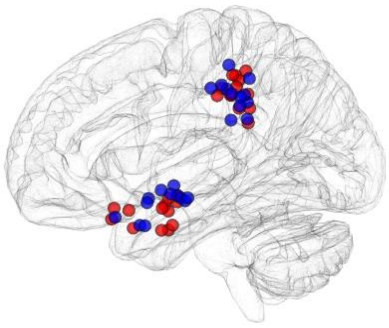

*Figure S4*. Functionally-localised targets for TMS shown in MNI space displayed on a glass brain (in Data Viewer 3D, Gouws et al., 2009). Red spheres = high semantic reliance participants; blue spheres = low semantic reliance participants.

The omnibus linear mixed effects model outcomes for behavioural data in the ISR task are shown in Table S3. Models for ISR data run separately by site are shown for SMG in Table S4 and ATL in Table S5. The linear mixed effects model outcomes for behavioural data the pattern span task in Table S6.

Table S3. *Linear mixed effects model outcomes for ISR data using Group, Site, TMS, Stimulus type, and covariates*

| | Group | Site | Stimulus | TMS | Item position | Type III tests F-value (DF) | z-value | Type II tests p-value | Parameter estimate | 95% CI | -2 Log Likelihood |
|---|---|---|---|---|---|---|---|---|---|---|---|
| Empty Model | | | | | | | | | | | 22495.00 |
| Full model | | | | | | | | | | | 21187.84 |
| Fixed effects: | | | | | | | | | | | |
| Intercept | | | | | | | | | 3.66 | (2.74 – 4.58) | |
| Group (high v low SR) | High | | | | | 11.79 (1, 22) | | .0006 | -0.44 | (-0.86 – -0.016) | |
| Site (ATL v SMG) | | ATL | | | | 0.15 (1, 18563) | | .70 | 0.15 | (-0.041 – 0.35) | |
| Stimulus (word v nonword) | | | Nonword | | | 405.85 (1, 18563) | | <.0001 | -2.40 | (-2.71 – -2.10) | |
| TMS (pre v post) | | | | Post | | 7.22 (1, 18563) | | .0072 | -0.073 | (-0.27 – 0.12) | |
| Group × site | High | ATL | | | | 11.71 (1, 18563) | | .0006 | -0.46 | (-0.72 – -0.20) | |
| Group × stimulus | High | | Nonword | | | 0.43 (1, 18563) | | .51 | -0.24 | (-0.52 – 0.035) | |
| Group × TMS | High | | | Post | | 2.68 (1, 18563) | | .10 | -0.14 | (-0.40 – 0.12) | |
| Site × stimulus | | ATL | Nonword | | | 3.18 (1, 18563) | | .074 | -0.054 | (-0.34 – 0.23) | |
| Site × TMS | | ATL | | Post | | 0.26 (1, 18563) | | .61 | -0.11 | (-0.39 – 0.16) | |
| Stimulus × TMS | | | Nonword | Post | | 0.46 (1, 18563) | | .50 | -0.17 | (-0.45 – 0.12) | |
| Group × site × stimulus | High | ATL | Nonword | | | 0.61 (1, 18563) | | .43 | 0.39 | (0.0012 – 0.78) | |
| Group × site × TMS | High | ATL | | Post | | 0.11 (1, 18563) | | .74 | 0.33 | (-0.038 – 0.69) | |
| Group × stimulus × TMS | High | | Nonword | Post | | 1.73 (1, 18563) | | .19 | 0.46 | (0.077 – 0.85) | |
| Site × stimulus × TMS | | ATL | Nonword | Post | | 0.05 (1, 18563) | | .82 | 0.25 | (-0.15 – 0.65) | |
| Group × site × stimulus × TMS | High | ATL | Nonword | Post | | 4.01 (1, 18563) | | .045 | -0.56 | (-1.11 – -0.012) | |
| Item position in list | | | | | 1 | 888.21 (6, 18563) | | <.0001 | 1.77 | (1.39 – 2.16) | |
| | | | | | 2 | | | | 1.10 | (0.71 – 1.48) | |
| | | | | | 3 | | | | 0.46 | (0.075 – 0.84) | |
| | | | | | 4 | | | | -0.0017 | (-0.38 – 0.38) | |
| | | | | | 5 | | | | -0.62 | (-1.0024 – -0.24) | |
| | | | | | 6 | | | | -0.33 | (-0.71 – 0.045) | |
| List size | | | | | | 71.99 (1, 18563) | | <.0001 | -0.50 | (-0.61 – -0.38) | |
| Random effects: | | | | | | | | | | | |
| Subject variance | | | | | | | 3.32 | .0005 | 0.19 | | |
| Item variance | | | | | | | 12.47 | <.0001 | 0.80 | | |

Table S4.

*Linear mixed effects model outcomes for ISR data, restricted to SMG, using Group, Stimulus type, TMS and covariates*

| | Group | Stimulus | TMS | Item position | Type III tests F-value (DF) | z-value | Type III tests p-value | Parameter estimate | 95% CI | -2 Log Likelihood |
|---|---|---|---|---|---|---|---|---|---|---|
| Empty Model | | | | | | | | | | 11651.43 |
| Full model | | | | | | | | | | 10837.84 |
| Fixed effects: | | | | | | | | | | |
| Intercept | | | | | | | | 4.73 | (3.46 – 5.10) | |
| Group (high v low SR) | High | | | | 6.92 (1, 22) | | .015 | -0.45 | (-0.90 – 0.00071) | |
| Stimulus (word v nonword) | | Nonword | | | 304.82 (1, 8979) | | <.0001 | -2.56 | (-2.90 – -2.21) | |
| Group × stimulus | High | Nonword | | | 0.02 (1, 8979) | | .90 | -0.25 | (-0.52 – 0.027) | |
| TMS (pre v post) | | | Post | | 4.62 (1, 8979) | | .03 | -0.061 | (-0.25 – 0.13) | |
| Group × TMS | High | | Post | | 0.75 (1, 8979) | | .38 | -0.15 | (-0.41 – 0.11) | |
| Stimulus × TMS | | Nonword | Post | | 0.36 (1, 8979) | | .55 | -0.18 | (-0.46 – 0.11) | |
| Group × stimulus × TMS | High | Nonword | Post | | 5.67 (1, 8979) | | .02 | 0.47 | (0.083 – 0.86) | |
| Item position in list | | | | 1 | 79.86 (1, 8979) | | <.0001 | 1.38 | (0.83 – 1.93) | |
| | | | | 2 | | | | 0.79 | (0.24 – 1.34) | |
| | | | | 3 | | | | 0.10 | (-0.44 – 0.65) | |
| | | | | 4 | | | | -0.31 | (-0.85 – 0.24) | |
| | | | | 5 | | | | -0.91 | (-1.45 – -0.37) | |
| | | | | 6 | | | | -0.57 | (-1.12 – -0.017) | |
| List size | | | | | 57.18 (1,8979) | | <.0001 | -0.63 | (-0.79 – -0.47) | |
| Random effects: | | | | | | | | | | |
| Subject variance | | | | | | 3.19 | .0007 | 0.23 | | |
| Item variance | | | | | | 9.87 | <.0001 | 0.69 | | |

Table S5.

*Linear mixed effects model outcomes for ISR data, restricted to ATL, using Group, Stimulus, TMS and covariates*

| | Group | Stimulus | TMS | Item position | Type III tests F-value (DF) | z-value | Type III tests p-value | Parameter estimate | 95% CI | -2 Log Likelihood |
|---|---|---|---|---|---|---|---|---|---|---|
| Empty Model | | | | | | | | | | 11547.71 |
| Full model | | | | | | | | | | 10781.02 |
| Fixed effects: | | | | | | | | | | |
| Intercept | | | | | | | | 2.83 | (1.59 – 4.067) | |
| Group (high v low SR) | High | | | | 15.22 (1, 22) | | .0008 | -0.87 | (-1.29 – -0.44) | |
| Stimulus (word v nonword) | | Nonword | | | 213.96 (1, 8973) | | <.0001 | -2.30 | (-2.65 – -1.95) | |
| Group × stimulus | High | Nonword | | | 0.86 (1, 8973) | | .35 | 0.14 | (-0.14 – 0.41) | |
| TMS (pre v post) | | | Post | | 2.64 (1, 8973) | | .10 | -0.19 | (-0.38 – 0.0080) | |
| Group × TMS | High | | Post | | 1.87 (1, 8973) | | .17 | 0.18 | (-0.082 – 0.44) | |
| Stimulus × TMS | | Nonword | Post | | 0.14 (1, 8973) | | .71 | 0.08 | (-0.21 – 0.37) | |
| Group × stimulus × TMS | High | Nonword | Post | | 0.19 (1, 8973) | | .66 | -0.086 | (-0.48 – 0.30) | |
| Item position in list | | | | 1 | 84.89 (1, 8973) | | <.0001 | 2.034 | (1.48 – 2.59) | |
| | | | | 2 | | | | 1.30 | (0.75 – 1.85) | |
| | | | | 3 | | | | 0.75 | (0.20 – 1.29) | |
| | | | | 4 | | | | 0.29 | (-0.25 – 0.84) | |
| | | | | 5 | | | | -0.42 | (-0.97 – 0.13) | |
| | | | | 6 | | | | -0.17 | (-0.71 – 0.38) | |
| List size | | | | | 20.48 (1,8973) | | <.0001 | -0.37 | (-0.53 – -0.21) | |
| Random effects: | | | | | | | | | | |
| Subject variance | | | | | | 3.19 | .0007 | 0.19 | | |
| Item variance | | | | | | 10.31 | <.0001 | 0.80 | | |

Table S6.

*Linear mixed effects model outcomes for pattern task data (TMS control task) using Group, Site, TMS, and covariates*

| | Group | Site | TMS | Item position | Type III tests F-value (DF) | z-value | Type III tests p-value | Parameter estimate | 95% CI | -2 Log Likelihood |
|---|---|---|---|---|---|---|---|---|---|---|
| Empty Model | | | | | | | | | | 1275.05 |
| Full model | | | | | | | | | | 1246.82 |
| Fixed effects: | | | | | | | | | | |
| Intercept | | | | | | | | 1.66 | (-0.081 – 3.40) | |
| Group (high v low SR) | High | | | | 0.58 (1, 21) | | .46 | -0.29 | (-1.052 – 0.47) | |
| Site (ATL v SMG) | | ATL | | | 0.69 (1, 921) | | .41 | 0.040 | (-0.51 – 0.59) | |
| Group × site | High | ATL | | | 0.03 (1, 921) | | .86 | -0.0041 | (-0.77 – 0.76) | |
| TMS (pre v post) | | | Pre | | 1.58 (1, 921) | | .21 | -0.35 | (-0.90 – 0.20) | |
| Group × TMS | High | | Pre | | 0.36 (1, 921) | | .55 | 0.21 | (-0.55 – 0.97) | |
| Site × TMS | | ATL | Pre | | 0.31 (1, 921) | | .58 | 0.20 | (-0.58 – 0.97) | |
| Group × site × TMS | High | ATL | Pre | | 0.02 (1, 921) | | .88 | -0.086 | (-1.16 – 0.99) | |
| Block order | | | | 1 | 1.93 (1, 921) | | .045 | 0.57 | (-0.039 – 1.18) | |
| | | | | 2 | | | | 0.48 | (-0.13 – 1.086) | |
| | | | | 3 | | | | 1.086 | (0.47 – 1.70) | |
| | | | | 4 | | | | 0.48 | (-0.13 – 1.086) | |
| | | | | 5 | | | | 0.62 | (0.0074 – 1.22) | |
| | | | | 6 | | | | 0.71 | (0.010 – 1.32) | |
| | | | | 7 | | | | 0.62 | (0.0074 – 1.22) | |
| | | | | 8 | | | | 0.94 | (0.33 – 1.56) | |
| | | | | 9 | | | | 0.29 | (-0.32 – 0.90) | |
| Array size | | | | | 7.38 (1,921) | | .0067 | -0.15 | (-0.26 – -0.042) | |
| Random effects: | | | | | | | | | | |
| Subject variance | | | | | | 2.46 | .0069 | 0.34 | | |

### Follow-up bootstrapped TMS ISR analyses

To help mitigate concern about the strength of the TMS evidence given the small groups and TMS effect sizes, we ran one-tailed paired-samples t-tests with bootstrapping (2,000 samples, bias corrected), in model-predicted recall accuracy following TMS (pre vs post-TMS) as a function of stimulation site and list type separately for high and low SR participants, to verify whether the significant patterns of disruption held. They did: These analyses confirmed significant disruption of nonword recall by SMG-TMS for the low-SR group only (nonwords: low SR group pre-TMS $M$ = .47, $SD$ = .10; post-TMS $M$ = .42 $SD$ = .13), $t(10)$ = 1.92, $p$ = .042 (one-tailed) BCa CI [.002, .08]. Whereas there was no SMG-TMS disruption of predicted nonword recall in the high-SR group (nonwords high-SR group pre-TMS $M$ = .36, $SD$ = .15; post-TMS $M$ = .37, SD = .13), $t(12)$ = -.50, $p$ = .31 (one-tailed) BCa CI [-.06, .03]). Bootstrapping analyses likewise supported SMG-TMS significantly disrupting word recall in the high-SR group only (words: low SR group pre-TMS *M* = .71, *SD* = .10; post-TMS *M* = .70, *SD* = .13; *t*(12) = 1.91, *p* = .04 [.003, .07]; high-SR group pre-TMS *M* = .66, *SD* = .11; post-TMS *M* = .63, *SD* = .12), *t*(10) = .57, *p* = .29 [-.03, .06]).

For completeness, the equivalent bootstrapping analyses of ATL-TMS indicated no significant disruption of recall, with only a non-significant trend for reduced word recall in the low-SR group (low-SR words pre-TMS *M* = .74, *SD* = .07; post-TMS *M* = .71, *SD* = .12, *t*(10) = 1.42, *p* = .09 (one-tailed) BCa CI [-.01, .06]; low-SR nonwords pre-TMS *M* = .48, *SD* = .07; post-TMS *M* = .46, *SD* = .11, *t*(10) = 0.89, *p* = .20 (one-tailed) BCa CI [-.02, .06]; high-SR words pre-TMS M = .61, SD = .12; post-TMS *M* = .61 *SD* = .12, t(12) = -.04, p = 0.49 (one-tailed) BCa CI [-.05, .04]; high-SR nonwords pre-TMS *M* = .36, *SD* = .12; post-TMS *M* = .36, *SD* = .13, *t*(12) = -.01, *p* = .50 (one-tailed) BCa CI [-.04, .03]).

**Supplemental material references**

Gouws A, Woods W, Millman R, Morland A, Green G (2009) Dataviewer3D: An open-source, cross-platform multi-modal neuroimaging data visualization tool. Front Neuroinform 3.

Savill NJ, Cornelissen P, Pahor A, Jefferies E (2019) rTMS evidence for a dissociation in short-term memory for spoken words and nonwords. Cortex 112:5–22.

Vatansever D, Bzdok D, Wang H, Mollo G, Sormaz M, Murphy C, Karapanagiotidis T, Smallwood J, Jefferies E (2017) Varieties of semantic cognition revealed through simultaneous decomposition of intrinsic brain connectivity and behaviour. Neuroimage 158:1–11 Available at: http://dx.doi.org/10.1016/j.neuroimage.2017.06.067.

Zhang M, Savill N, Margulies DS, Smallwood J, Jefferies E (2019) Distinct individual differences in default mode network connectivity relate to off-task thought and text memory during reading. Sci Rep 9.
